# Substrate stiffness facilitates improved induced pluripotent stem cell production through modulation of both early and late phases of cell reprogramming

**DOI:** 10.1101/2023.02.27.530366

**Authors:** Mohammad Mahfuz Chowdhury, Samuel Zimmerman, Hannah Leeson, Christian Maximilian Nefzger, Jessica Cara Mar, Andrew Laslett, Jose Maria Polo, Ernst Wolvetang, Justin John Cooper-White

## Abstract

Cell reprogramming involves time-intensive, costly processes that ultimately produce low numbers of reprogrammed cells of variable quality. By screening a range of polyacrylamide hydrogels (pAAm gels) of varying stiffness (1 kPA – 1.3 MPa) we found that a gel of medium stiffness significantly increases the overall number of reprogrammed cells by up to ten-fold with accelerated reprogramming kinetics, as compared to the standard Tissue Culture PolyStyrene (TCPS)-based protocol. We observe that though the gel improves both early and late phases of reprogramming, improvement in the late (reprogramming prone population maturation) phase is more pronounced and produces iPSCs having different characteristics and lower remnant transgene expression than those produced on TCPS. Comparative RNA-Seq analyses coupled with experimental validation reveals that modulation of Bone Morphogenic Protein (BMP) signalling by a novel reprogramming regulator, Phactr3, upregulated in the gel at an earliest time-point without the influence of transcription factors used for reprogramming, plays a crucial role in the improvement in the early reprogramming kinetics and overall reprogramming outcomes. This study provides new insights into the mechanism via which substrate stiffness modulates reprogramming kinetics and iPSC quality outcomes, opening new avenues for producing higher numbers of quality iPSCs or other reprogrammed cells at shorter timescales.

## 1. Introduction

The ability to reprogram somatic cells (e.g., skin fibroblasts) to induced pluripotent stem cells (iPSCs) and derive unlimited numbers of all types of somatic cells (with or without gene editing) from them has created exciting new pathways for developing better disease models for efficient drug screening or achieving patient-specific treatments using cell therapies.[1, 2]. However, deriving safe and high-quality iPSC (in terms of tumorigenic and immunogenic potential, genomic, transcriptomic, and epigenetic stability, and variability in differentiation capacity) is still a lengthy and costly endeavour, representing a major challenge in transitioning iPSC research into clinical or drug screening applications.[3–5]

Somatic cell reprogramming to iPSCs can be divided into three phases/stages: the early or initiation stage, the duration of which is 3 days to less than a week and characterised by a mesenchymal-to-epithelial transition; the late/maturation stage, from the end of early phase to the end of reprogramming culture; and the stabilization stage, which covers early through to late passaging of iPSCs.[6] Whilst any improvement at the early-stage of reprogramming (entry of somatic cells to reprogramming-prone cells) represents a favourable outcome, improvement in the late stage of reprogramming is crucial, as many reprogramming-prone cells from the early stage fail to establish their pluripotency gene network in the late or maturation phase, thereby reducing the overall reprogramming efficiency.[6] It has been suggested that increasing the kinetics and efficiency of the reprogramming process would also likely positively impact iPSC quality/characteristics and reduce the arduous and expensive tasks of screening for high-quality iPSCs. [3]

It is clear that the original somatic cell quality exerts influence over the derived iPSC product[7], and that the reprogramming methodologies and culture conditions used to reprogram cells also influence reprogramming efficiency and iPSC quality.[3, 4] Reprogramming efficiency has also been shown to be enhanced by a variety of biochemical-based methods (e.g., mRNA, protein or small molecules)[8, 9], however it remains largely unclear whether, and how, biophysical cues presented to cells can improve reprogramming outcomes. Matrix/substrate ‘stiffness’ has been shown to be a key regulator of mechanotransduction pathways impacting adult and pluripotent stem cell behaviour, but to date our understanding of the impact of substrate stiffness on cell reprogramming outcomes remains limited.[10–19] Choi and co-workers[17] successfully reprogrammed mouse fibroblasts by plating them onto 2D polyacrylamide hydrogel substrates of varying stiffness (ranging from 0.1 kPa to 20 kPa) for one day, then replating them on TCPS for the remainder of the reprogramming process. They observed 2-times more miPSCs generated on TCPS that was inoculated with day 1 cells from the 0.1 kPa substrate, compared to those from the 20 kPa substrate. By constructing polyethylene glycol-based 3D hydrogels of a very small range of stiffnesses (from 0.3 to 1.2 kPa), Caiazzo and co-workers[19] reported that the physical confinement exerted by the 3D hydrogel on cells induced their earlier entry into MET (Mesenchymal-to-Epithelial Transition) and epigenetic modification at the early stage of reprogramming that ultimately produced 2.5- and < 2- times more mouse and human iPSC colonies, respectively,[11] compared to standard 2D controls.

Most recently, Kim and co-workers[20] demonstrated improvements of mouse fibroblast reprogramming to iPSC by introducing already transduced fibroblasts (after 3 days on TCPS) into a 3D methacrylated hyaluronic acid-based hydrogel, which had a stiffness of 0.15 kPa. These previous studies suggest that hydrogels of very low stiffness improved reprogramming outcomes through the modulation in known mechanosensitive (e.g., YAP/TAZ) signalling pathways and/or epigenetic changes impacting MET at the early stage of reprogramming.

The substrates investigated to date have been significantly lower (by multiple orders of magnitude) in stiffness than TCPS (i.e., ≥ 10^6^ kPa)[21]. Moreover, due to the way in which previous investigations into the impact of substrate stiffness on reprogramming have been performed (as highlighted above), there is little or no information available on the kinetics of the reprogramming process from start to end on substrates of varying stiffness (as no study has achieved such an outcome), and whether these substrates produce iPSC of different characteristics to those produced from TCPS. Such information could be crucial if wishing to combine other biomaterial features (e.g., topography[18]) and soluble factors or small molecules with these substrates to optimise the reprogramming process for producing quality iPSCs at shorter timescale and less cost, facilitating their effective use in regenerative medicine or drug screening.

In this study, we have systematically assessed the extent to which hydrogel-based culture substrates of varying stiffness over ranges of 0.1 kPa to 1300 kPa modulate the transition of murine and human somatic fibroblasts to iPSCs. Unlike most of the protocols used in investigating reprogramming on different biomaterial substrates[17,18,20], we herein established a consistent gel condition for continuous reprogramming of both human and mouse fibroblasts to iPSCs that did not require fibroblast replating at any time during the reprogramming process. We show how the gel condition impacted the early and the late stages of the reprogramming and overall reprogramming outcomes. Furthermore, by combining reprogramming time-course RNA-Seq analysis and experimental observations, we evaluated reprogramming kinetics and the characteristics of iPSCs produced under the gel and TCPS conditions and identified a novel and critical regulator of the observed improvements.

## 2. Results

### 2.1 An optima exists for the stiffness of hydrogels for MEF reprogramming

To systematically investigate the role of substrate stiffness on cell reprogramming we seeded murine embryonic fibroblasts (MEFs) with doxycycline-inducible OKSM reprogramming factors and a OCT4-GFP reporter on polyacrylamide (pAAm) gels of varying elasticity (*E* from 1 kPa to 1.3 MPa, **Figure S1**, Supporting Information) with Gelatin (0.2% w/v) that was immobilised on the surface with the Sulfo-SANPAH (S-S) conjugation method and assessed reprogramming efficiency into miPSCs by enumerating GFP expressing colonies. This revealed that across the range of gels examined, MEF reprogramming on 102 kPa and 247 kPa gels (**Figure S2a**, Supporting Information) produced four and three-fold higher numbers of Oct4 GFP+ miPSC colonies compared to Gelatin coated TCPS.

However, these S-S treated gels were unable to support long term culture, as indicated by frequent detachment of cells from the culture substrate (**Figure S2b**, Supporting Information). We hypothesized that this was due to suboptimal immobilization of Gelatin on these pAAm substrates with the S-S method.[22], leading us to examine an alternate conjugation method, using L-DOPA.[23]

Satisfyingly, the L-DOPA conjugation method produced a more homogenous extracellular matrix (ECM) coating than that achieved with S-S treatment (**Figure S3**, Supporting Information) and we confirmed that equal amounts of Gelatin were deposited on the pAAm gels of varying stiffness and TCPS treated with L-DOPA (**Figure S3**, Supporting Information). When MEFs were reprogrammed on L-DOPA-treated pAAm gels with range of moduli, we found that pAAm gels of *E* = 102 kPa and 1.3 MPa consistently produced four-fold more Oct4 GFP+ colonies than the TCPS control (**Figure 1a, b**). In contrast, pAAm gels of 1 kPa and 16 kPa showed very low numbers of Oct4 GFP+ colonies compared to the TCPS control (**Figure 1a, b**).

**Figure 1.**
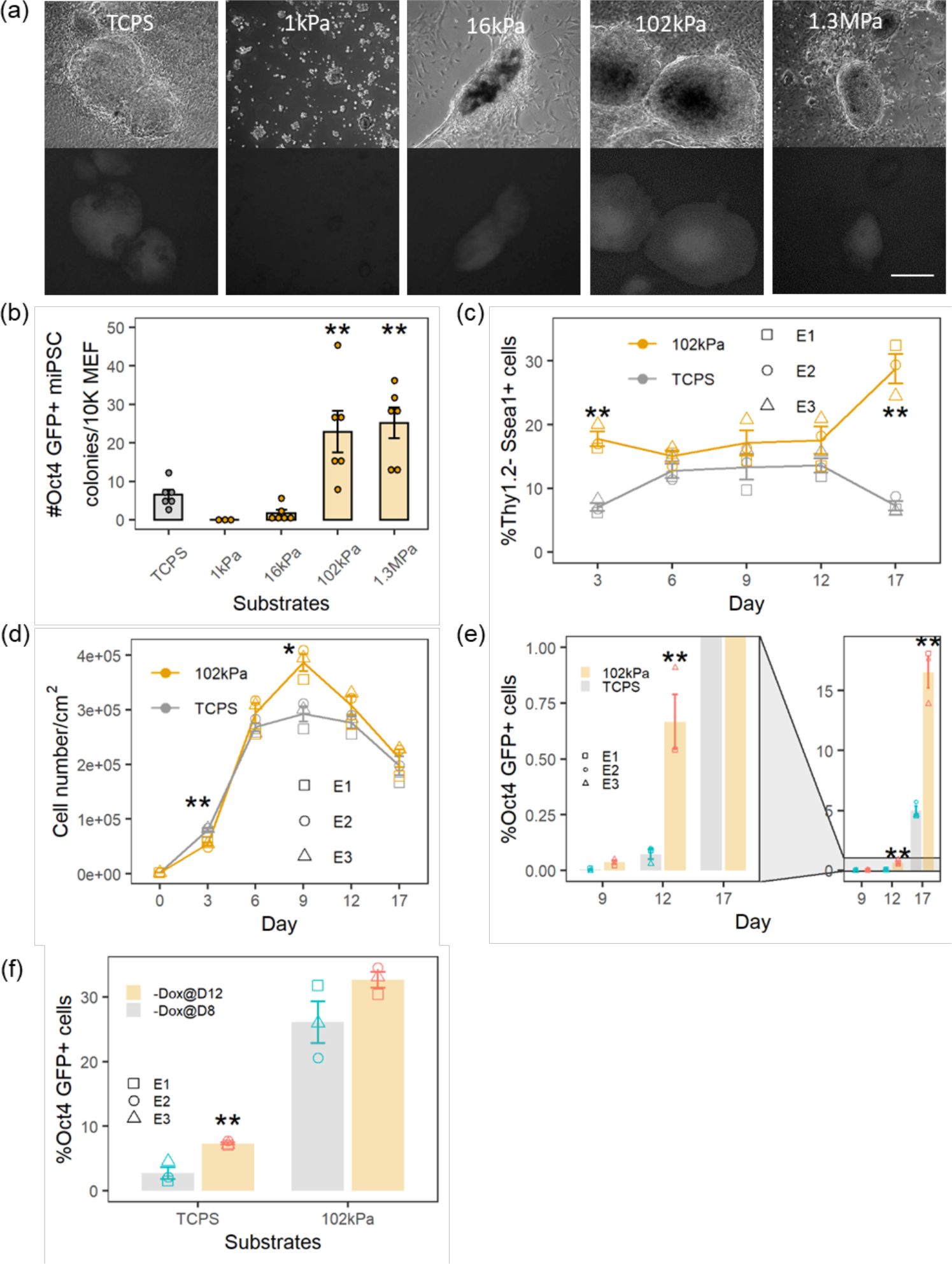
MEF reprogramming on pAAm gels of various stiffness, which were conjugated with L-DOPA for ECM immobilization. (**a**) Brightfield (top row) and corresponding GFP channel (bottom row) showing miPSC colonies on TCPS and pAAm gels. (**b**) Number of Oct4 GFP+ miPSC colonies counted at day 17 on TCPS and pAAm gel of various stiffness. (**c**) Percentage of cells undergoing reprogramming (Thy1.2- SSEA1+ cells) on TCPS and pAAm gel of 102 kPa at various days. (**d**) Cell growth, as quantified by cell number on each day, over the reprogramming period in TCPS and pAAm. (**e**) Emergence of Oct4 GFP+ in reprogramming culture on TCPS and pAAm over time starting from Day 9. (**f**) % of Oct4 GFP+ miPSCs at day 17 on TCPS and pAAm gel after Doxycycline was withdrawn earlier (Day 8) than the normal (Day 12). In all cases, MEF from three different embryos and at least two technical replicates were used. Data is represented mean ± SEM and “***”, “**” and “*” represents *p* < 0.001, *p* < 0.01 and *p* < 0.05, respectively. Scale bar represents 500 µm.

Since the reprogramming of MEFs into iPSC consists of three separate stages characterised by an initial downregulation of the fibroblast-associated marker Thy1.2 (day 1–3), activation of the SSEA1 antigen (day 3–9) and eventually upregulation of the Oct4-GFP reporter (day 9–17)[24], we were interested to find out which phase of the reprogramming process was affected by substrate stiffness. Since cell reprogramming is intimately connected to cell cycle progression, we first examined cell proliferation on the pAAm gels of varying stiffness, revealing that cell proliferation on all pAAm gels by D3 was reduced as compared to TCPS controls (**Figure S4a**, Supporting Information).

Nevertheless, all pAAm gels produced higher numbers of Thy1.2-/SSEA-1+ cells compared to the control TCPS on Day 3 (D3) of reprogramming (**Figure S4b**, Supporting Information).

However, pAAm gels with *E* below 102 kPa were unable to produce Oct4 expressing iPSC (**Figure 1b**), suggesting that pAAm substrates with *E* below 102 kPa did not support later stages of reprogramming. Since regardless of the conjugation method used, 102 kPa gel consistently produced higher numbers of iPSC colonies compared to the TCPS, we focussed on examining the cell reprogramming process on this gel in detail.

At D3, the %Thy1.2-/SSEA-1+ cell population was three-fold higher on the 102 kPa pAAm gel compared to TCPS control (**Figure 1c**). Between D6 to D12, the number of Thy1.2-/SSEA-1+ cells on either the 102 kPa gel or TCPS did not substantially change. However, from D12 to D17, we observed a robust increase in the number of Thy1.2-/SSEA-1+ cells on pAAm as compared to TCPS, which showed a decrease (**Figure 1c**) in Thy1.2-/SSEA-1+ cells. Though cell growth was noted to decrease on pAAm by D3, it gradually increased during D3 to D9 and maintained a higher cell growth trend (non-significant) over the remaining reprogramming period (**Figure 1d**).

Between day 9 and day 17, the 102 kPa gels fostered significant increases in the number of Oct4-GFP+ cells as compared to TCPS (**Figure 1e**), indicating substrate stiffness accelerated the transition of Thy1.2-/SSEA-1+ cell into pluripotent Thy1.2-/SSEA-1+/Oct4 GFP+ miPSC cells. As expected,[25] the removal of Doxycycline (Dox) on D8 from cells cultured on TCPS resulted in a significant reduction in GFP+ cells (**Figure 1f**). In contrast, removal of Doxycycline (Dox) on D8 from cells reprogrammed on the 102 kPa pAAm gels condition did not affect the number of GFP+ cells, suggesting that the mechanics of the hydrogel substrate enhanced entry into pluripotency at the early stage, and can independently drive the improvement of reprogramming, irrespective of the modulation of reprogramming at the late phase.

Overall, these data indicated that reprogramming of MEFs on a pAAm hydrogel substrate of ∼102 kPa leads to a four-fold increase in Oct4 GFP+ miPSCs compared to TCPS by enhancing the number of Thy1.2-/SSEA-1+ cells during the initial phase of reprogramming (D0 to D3) and by fostering the maturation of the pluripotent state during the final reprogramming phase (D9 to D17).

### 2.2 Human fibroblast reprogramming on optimised hydrogel substrate

We next wished to assess whether 102 kPa pAAm gels would also enhance Sendai virus- mediated reprogramming of human neonatal dermal fibroblasts (hDFn). To avoid the replating of cells, a process that is customary in the standard Sendai virus protocol on TCPS, we used lower fibroblast seeding densities than usual, allowing us to assess the effect of substrate stiffness on the entire human fibroblast population during the 18-day reprogramming process.

The 102 kPa pAAm gel condition produced two-times higher ALP+ hiPSC colonies compared to TCPS (**Figure 2a, b** and Supporting Information **Figure S5a**) at D18, demonstrating for the first time the efficacy of 102 kPa pAAm gel in improving human cell reprogramming. Time- course profiles of human cells undergoing reprogramming revealed an increase in CD13- SSEA4+Tra1-60+/- cells at earlier time points (D4), relatively little increases during the middle period (D8 to D12) and a large increase in fully reprogrammed cells CD13-SSEA4+Tra1-60+ cells during the late reprogramming period (D12 to D18) (**Figure 2c, d**). The 102 kPa gel condition produced a modest increase (two times) in the reprogramming-prone (SSEA4+ Tra1-60+/-) populations compared to TCPS but produced six-times higher amounts of fully reprogrammed hiPSCs (CD13-SSEA4+Tra1-60+) at D18 (**Figure 2d**), again indicating that the 102 kPa hydrogel efficiently fostered entry and maturation of pluripotency, similar to what was observed with MEFs.

**Figure 2.**
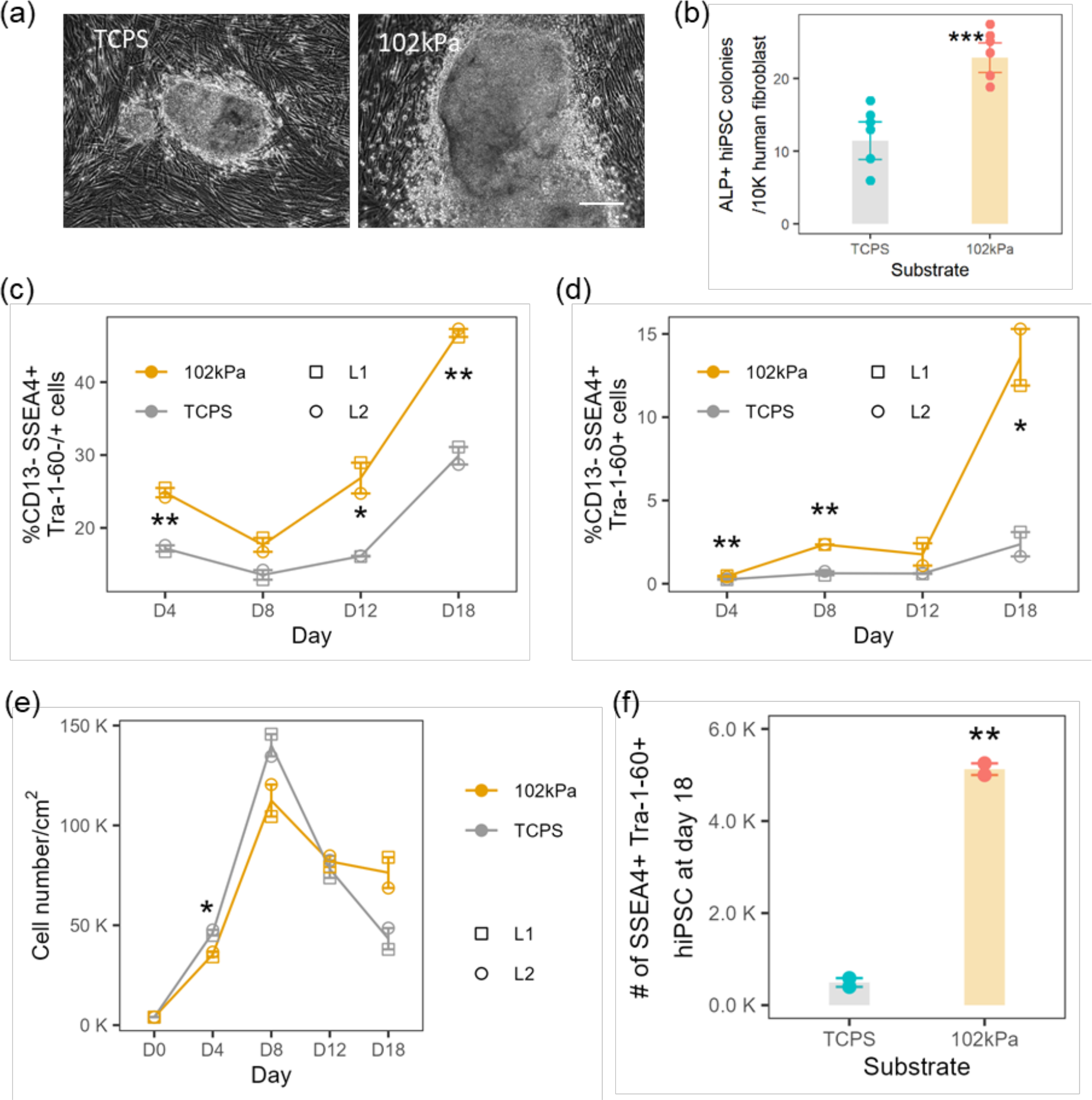
Comparison of neonatal human dermal fibroblasts (hDFn) reprogramming on 102 kPa pAAm gel and TCPS. **(a)** Bright field images of hiPSC colony and (b) number of ALP+ hiPSC colonies at day 18 in TCPS and pAAm gel. Percentage of **(c)** reprogramming prone CD13-SSEA4+Tra1-60-/+ cells and **(d)** bona fide reprogrammed SSEA4+Tra1-60+ cells over time in TCPS and pAAm gel. (**e**) cell growth (quantified by cell number) over time in TCPS and pAAm. (**f**) yield (number of iPSC per 10K inoculated hDFn at day 0) of hiPSC at day 18. For all data, hDFn were from two cell lines (L1 and L2) and a minimum of three technical replicates were used. Data is represented as mean ± SEM and “***”, “**” and “*” represents *p* < 0.001, *p* < 0.01 and *p* < 0.05, respectively. Scale bar represents 500µm.

Like in the case of MEF reprogramming, cell growth was slightly decreased at early period (D0 to D4) and drop in the cell growth at later stage (D8 to D18) was less prominent (**Figure 2e**) in pAAm gel compared to TCPS. At the end of reprogramming, cell number in the pAAm gel condition was two-times higher compared to the TCPS (**Figure 2e**). This increase, combined with the higher number of bona fide hiPSCs (SSEA4+ Tra1-60+), resulted in a *ten-times higher* yield of hiPSC from the pAAm gel compared to the TCPS (**Figure 2f**). These results demonstrated that, like MEF reprogramming, the pAAm gel condition accelerated human fibroblast reprogramming process by increasing the conversion of cells to pre-reprogrammed states during early stage and entry into a fully reprogrammed state during the final stage of reprogramming.

### 2.3 Hydrogel produced iPSCs with characteristics different from TCPS-produced iPSCs

To determine how the derivation of miPSC on 102 kPa gels impacted gene expression during reprogramming, we subjected triplicate samples at each time point to RNA-Seq and assessed total RNA expression in hydrogel and TCPS reprogrammed murine iPSCs following 5 passages on TCPS.

Bulk gene set enrichment analysis with differentially expressed genes (DEGs) at D17 between TCPS and pAAm miPSCs for various MIKKELSEN gene sets[26] in GSEA demonstrated that iPSC in pAAm upregulated dedifferentiated state gene sets and downregulated gene sets which are inactive in ESC and active in MEF (Supporting Information **Figure S6a**, see Bioinformatics analysis section in Methods for details) compared to the TCPS condition.

At an individual gene expression level, a heatmap (**Figure 3a**) of most varied DEGs at day 17 showed that mouse Oct4 GFP+ cells in the gel condition downregulated various markers that are highly expressed in primed mESC (Fgf5, T, Foxa2, Fgf8, Wnt8a and Cer1)[27–31]. Notably, these markers were in the top 25 genes that varied most between TCPS and pAAm conditions, indicating the primed status of miPSC was a significant factor discerning pAAm from TCPS. Moreover, miPSC on pAAm showed downregulation of Sox17, Lin28b and modest upregulation of Klf4, Tbx3 and Tfcp2l1 (Supporting information **Figure S6b and S7**), which are up- and down-regulated in primed mESC, respectively.[27,30,31]

**Figure 3.**
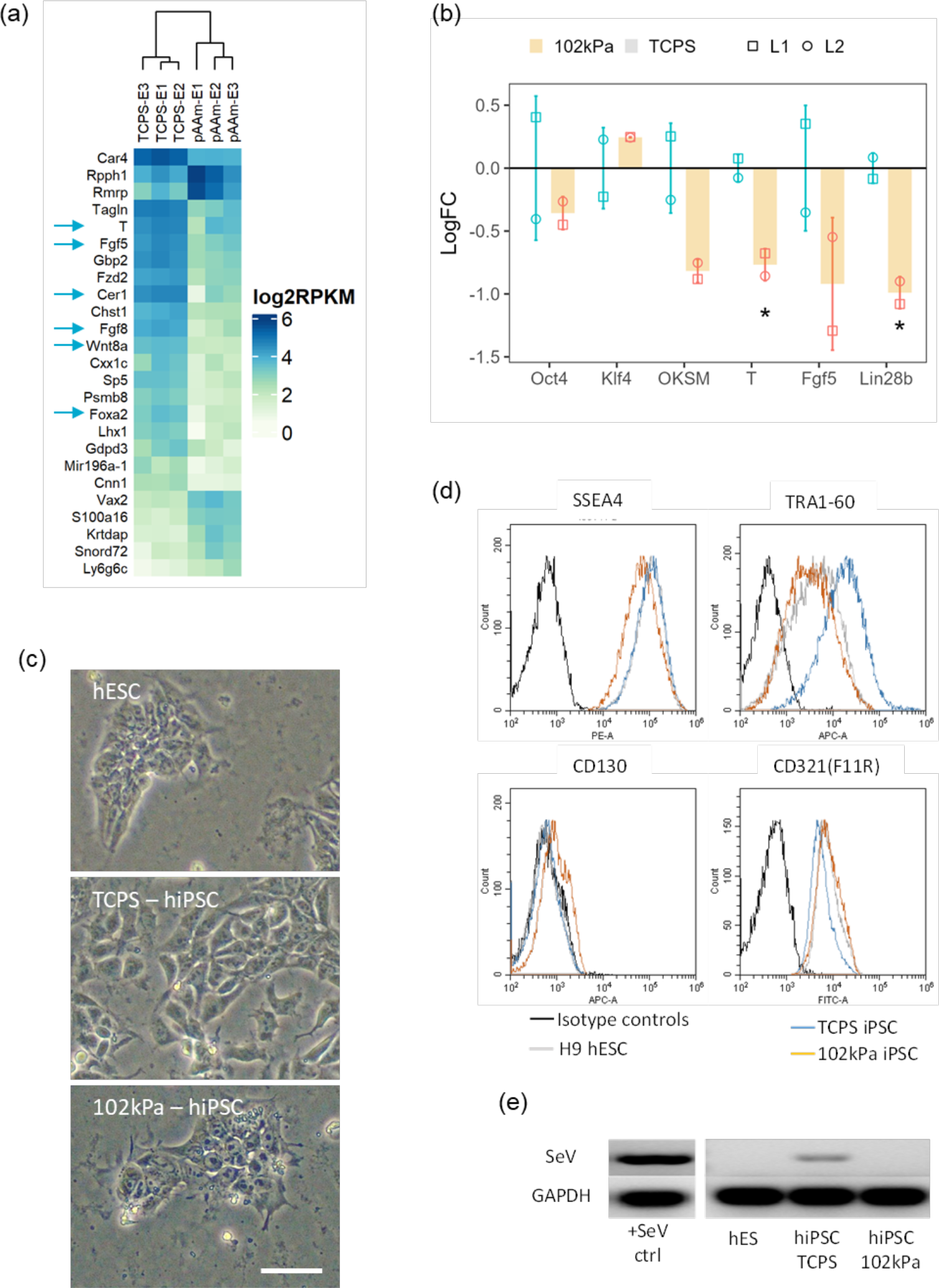
Characterization of mouse (**a** and **b**) and human (**c, d, and e**) iPSC produced on pAAm and TCPS. (**a**) Heatmap of most varied 25 DEGs which separated Oct4 GFP+ cell population at day 17 in TCPS and pAAm. Arrows indicate the genes that are discussed in the result text. (**b**) Expression level of common PSC (Oct4, Klf4) and primed markers (T, Fgf5 and Lin28b) and OKSM in miPSCs from pAAm and TCPS after subculturing them in TCPS for 5-times (n = 2). Line at y = 0 represents mean expression in TCPS. Error bar represents mean ± SD. (**c**) hiPSC morphology, (**d**) flow cytometric analyses of pluripotency associated cell- surface makers (that vary depending on hESC primed status), and (**e**) residual SeV expression quantified by RT-PCR and gel electrophoresis in hESC (human embryonic stem cell, H9), hiPSC from TCPS and pAAm gel condition (n = 1). hiPSCs from both pAAm and TCPS condition were subcultured for 8-times in TCPS condition before performing all the experiment. Scale bar represents 50 µm.

Next, we investigated whether the difference observed in passage 0 (P0) iPSC could also be observed after expansion of miPSCs from pAAm and TCPS under TCPS conditions. Interestingly, even after subculturing on TCPS for 5 times, qPCR analysis showed that miPSCs originally derived on the pAAm gel downregulated primed mESC markers (T and Lin28b) and showed lower expression trends for OKSM transgene and Fgf5 compared to the TCPS-derived miPSCs (**Figure 3b**). Overall, these observations indicated that pAAm produced miPSCs were characteristically different (i.e., presumably less primed) compared to the miPSCs produced on TCPS. Next, we wished to investigate whether these observations were also translated in the case of hiPSCs.

After expanding hiPSCs from both pAAm and TCPS under TCPS conditions for 8-times, hiPSCs from the gel condition were morphologically more like the H9 hESCs when compared to the TCPS-derived cells (**Figure 3c**) and exhibited a slower growth rate. Moreover, like the observed changes in expression pattern depending on the primed status of stem cells [32, 33], hiPSCs derived from pAAm showed lower expression of SSEA4 and Tra1-60 and higher F11R and CD130 compared to TCPS-derived cells (**Figure 3d**). We also observed that hiPSCs from pAAm almost lost all Sendai virus vector expression, whereas TCPS still had significant residual expression (**Figure 3e**). Overall, these results suggested that, like miPSCs, pAAm-produced hiPSCs are characteristically different when compared to TCPS-produced hiPSCs and that the reprogramming process on hydrogel facilitated faster removal of exogenous factors used for reprogramming.

Lastly, we confirmed that embryoid bodies generated from both miPSC and hiPSC sourced from TCPS and 102 kPa pAAm gel differentiated into cell types that expressed genes representative of each of the three germ layers, and hiPSC lines from both substrates were karyotypically normal (Supporting Information **Figure S8a-c**).

### 2.4 Hydrogel modulates signalling and metabolic pathways that support faster reprogramming kinetics and distinct iPSC characteristics

In a Principal Component Analysis (PCA, **Figure 4a**) of RNA-Seq data, day 3 and 6 reprogramming intermediates (Thy1-SSEA1+) from TCPS and pAAm gel clustered together, indicating similar kinetics during the early reprogramming period. However, intermediates in pAAm and TCPS at day 9 and day 12 did not cluster together and a large separation was observed between them at day 12. Moreover, compared to the TCPS, the separation between day 9 and 12 clusters (Thy1-SSEA1+ cells) and between day 12 and day 17 Oct4 GFP+ clusters were larger and smaller in case of pAAm gel, respectively, indicating faster progression of reprogramming intermediates to iPSC on the gel. These results also correlated well with the ability of pAAm gel to accelerate production of GFP+ iPSCs from an earlier time point compared to the TCPS **(Figure 1e**). Finally, at day17, Oct4 GFP+ bona fide iPSC generated from TCPS and pAAm gel clustered together.

**Figure 4.**
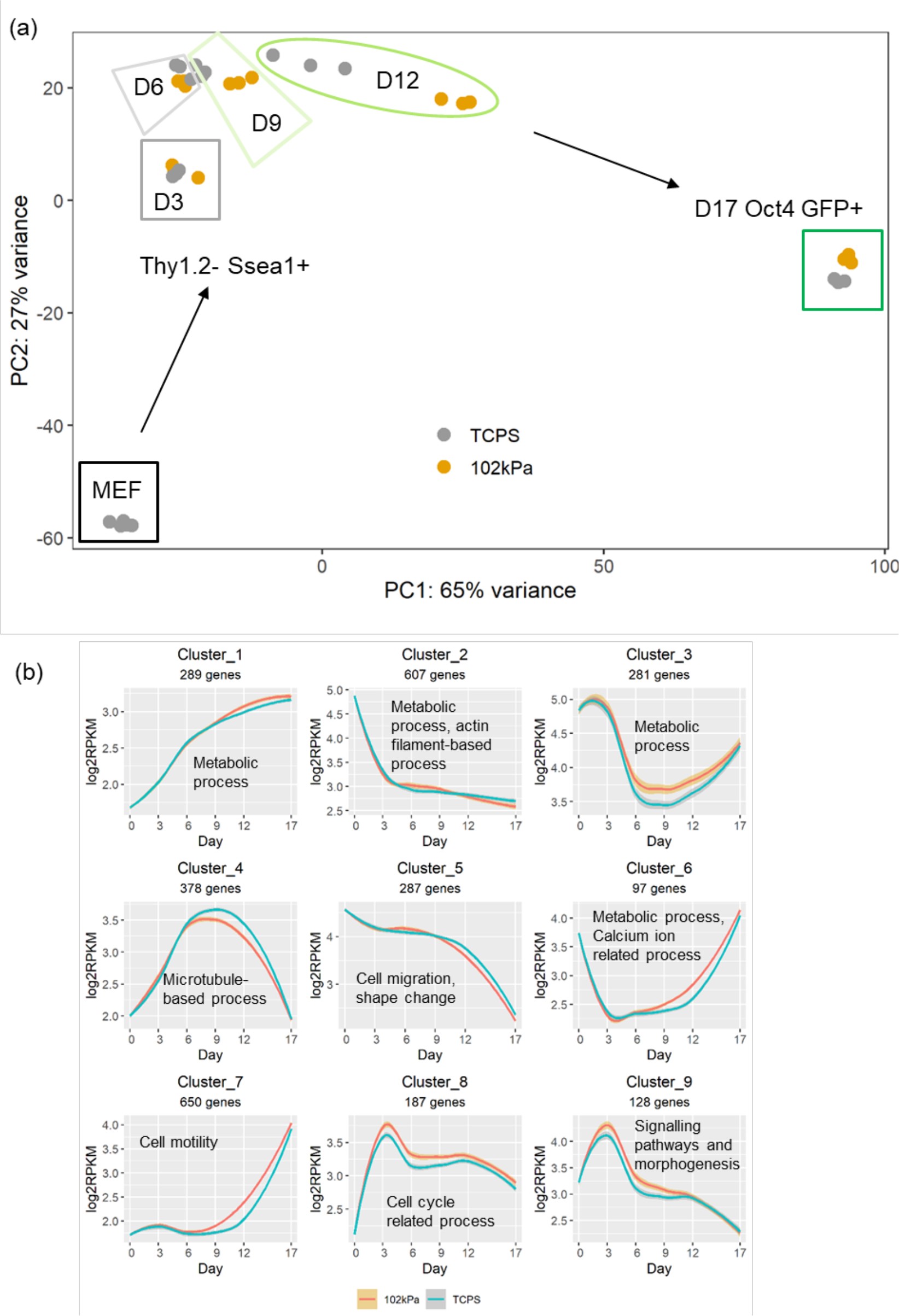
Characteristics of overall gene expression (quantified by RNA-Seq) change during reprogramming under different substrate condition. (**a**) Principal component analysis (PCA) showing progress of different populations over time. (**b**) DEGs (FDR = 0.01) between pAAm and TCPS at different time points were clustered hierarchically and contrasted with the corresponding expression in TCPS. Band enclosing cluster or expression profile in (b) represent 95% confidence interval. Text label in each plot representative of significantly regulated pathways obtained in Panther GO-SLIM analysis with the corresponding genes in a cluster.

PCA also indicated that reprogramming in pAAm and TCPS followed a similar reprogramming route. Clustering of DEGs (with FDR 0.01) between TCPS and pAAm at any time point within D3 to D12 and then profiling their temporal expression showed that profiles were similar in TCPS and pAAm, but that their expression levels differed at time periods (**Figure 4b**).

Collectively, these observations indicated that, rather than taking a different route through reprogramming, pAAm modulated cell reprogramming by more pronounced up or down regulation of genes.

Focal adhesion, ECM-receptor interaction, and Regulation of actin cytoskeleton, all of which characterise cell engagement with the underlying substrate, were down-regulated at day 3 but were upregulated from day 6 to day 12, at day 6, and from day 6 to day 9, respectively (**Figure 5**). Pathways regulating pluripotency of stem cells were up regulated in the pAAm condition from day 3 to day 12. At day 17, iPSCs on pAAm downregulated pathways related to cell adhesion (Focal adhesion, ECM-receptor interaction, and Cell adhesion molecules) as well as Axon guidance, and Antigen processing and presentation pathways compared to TCPS (**Figure 5**), which are reported to be downregulated in mESC and naïve hESC.[31,34,35]

**Figure 5.**
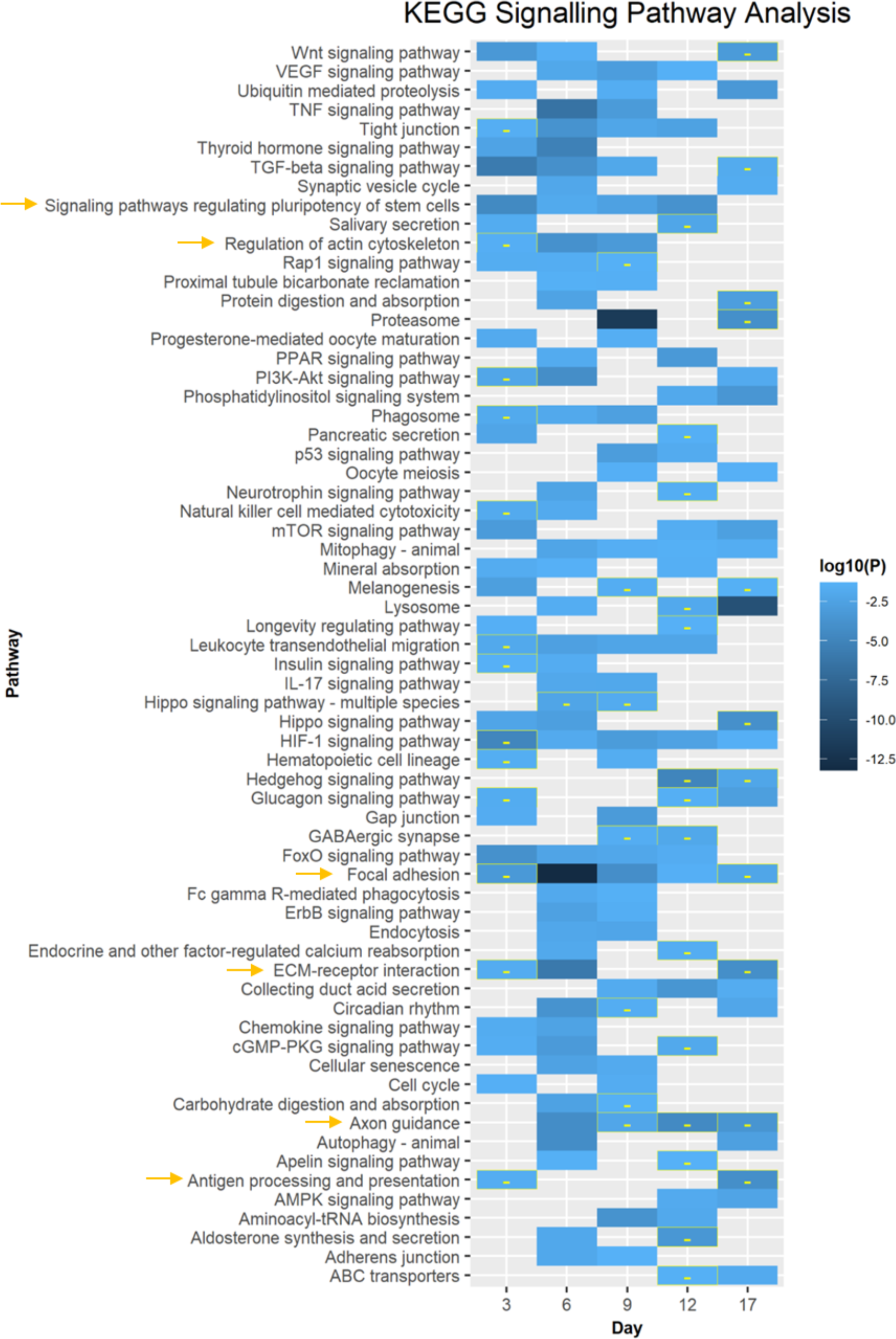
Up and Down regulated KEGG signaling pathways in pAAm at different time points compared to TCPS. All tiles indicate up regulation except “-” symbol containing tiles. Arrows indicate the pathways that were discussed in the text.

At day 3, reprogramming cells in pAAm gel significantly downregulated many metabolism pathways (which included Glycolysis/Gluconeogenesis, Citrate cycle and oxidative phosphorylation) compared to TCPS (**Figure 6**). After Day 3, cells on the pAAm hydrogel started to upregulate various metabolic pathways, which were maximised at the end of reprogramming.

**Figure 6.**
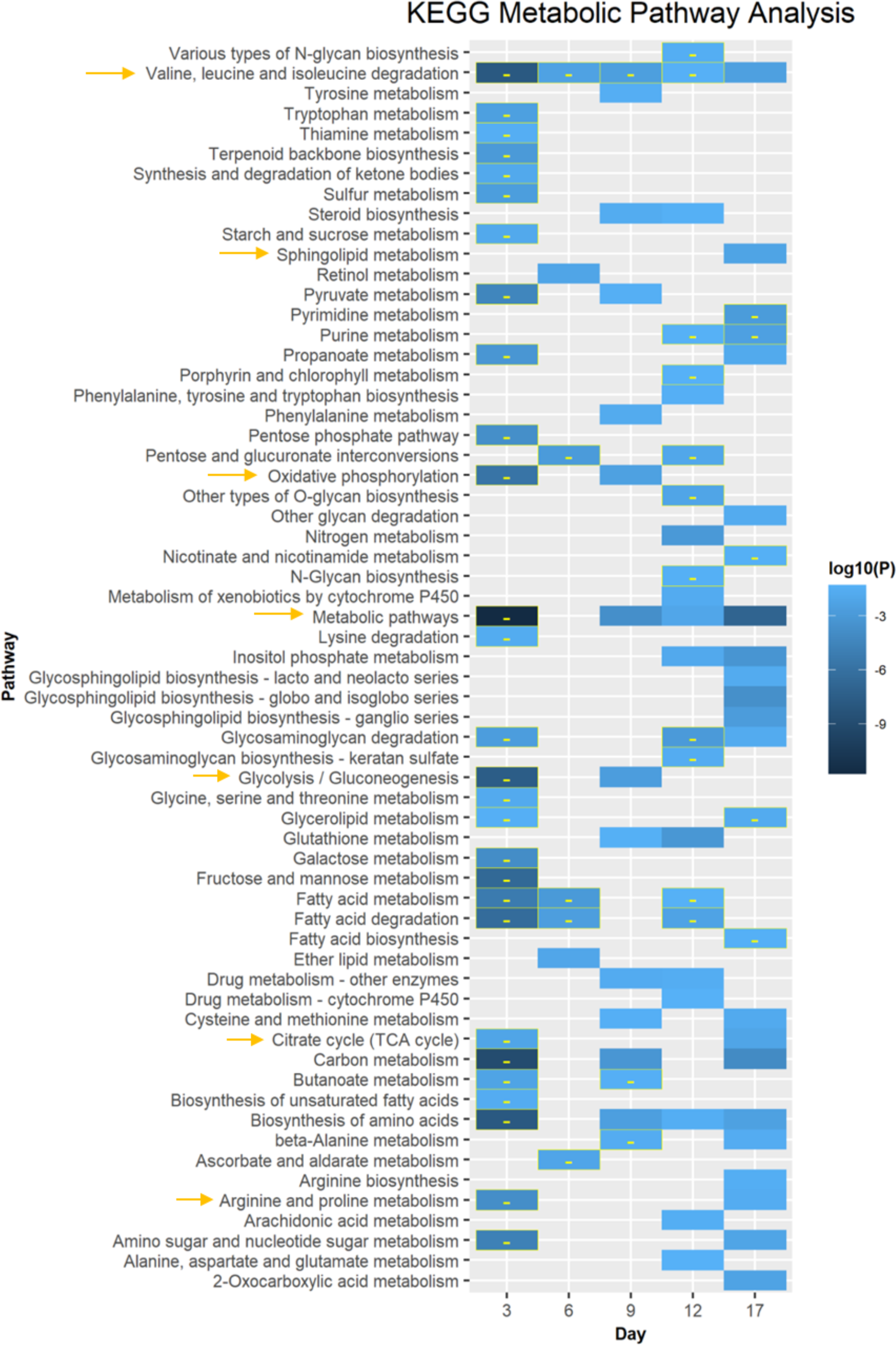
Up and Down regulated KEGG metabolic pathways in pAAm at different time points compared to TCPS. All tiles indicate up regulation except “-” symbol containing tiles. Arrows indicate the pathways that were discussed in the text.

Both glycolysis and oxidative phosphorylation (OxPhos) show rapid increase and decrease (forming a bell-shaped pulse) around day 3-4 of reprogramming before their gradual increase (glycolysis) and decrease (OxPhos) over the remaining period of reprogramming.[36] These data and faster kinetics of reprogramming in pAAm condition suggested that the down regulation of metabolic pathways at day 3 and upregulation thereafter might have represented a faster shift of metabolism in the pAAm condition compared to TCPS.

Although OxPhos and glycolysis/gluconeogenesis were similar in iPSCs generated on the pAAm gel and TCPS on day 17, Sphingolipid, various amino acid metabolism and TCA cycle were upregulated in the pAAm condition (**Figure 6**). Upregulation of the TCA cycle, along with downregulation of Lin28b in less primed pAAm iPSC, correlated well with the known Lin28b effect on PSC metabolism and their transition from naïve to primed.[30] Overall, regulation of both signalling and metabolic pathways in the pAAm condition are consistent with its supportive role in facilitating iPSC-production more efficiently compared to the TCPS condition.

### 2.5 Phactr3 critically impacted the early stage and end stage outcome of cell reprogramming

We next focused on identifying molecular targets which appear sensitive to substrate stiffness at the early stages of reprogramming for two reasons: entry into a reprogramming prone state at the early stage is a bottleneck for the overall reprogramming process and efficiency, and cell crowding and cell-remodelled ECM become additional important factors at the later stages of reprogramming, potentially overshadowing other substrate-induced impacts.

By investigating DEGs at D3, we identified various molecular pathways, including Bmp, Fgf and Gata transcription factors (**Figure S9a, S10**, Supporting Information; see Bioinformatics analysis section for details), which are known to accelerate cell reprogramming.[37–41] In terms of the Bmp signalling pathway, the pAAm condition upregulated various factors: Bmp2, Acvr1, Smads (4, 9), and Ids (1-3) (**Figure S9, S10**, Supporting Information). Hayashi and co-workers reported increased cell reprogramming efficiency by Ids (1-3), in which upregulation is mediated by Bmp/Acvr1 signaling. Bmp signalling contributes to established signalling pathways regulating pluripotency of stem cells and is crucial for MET and successful reprogramming in conventional TCPS culture conditions [37, 42]. Choi et al. suggested improvement in cell reprogramming by short-term hydrogel exposure were similarly due to MET.[17] Overall, these results on the 102 kPa substrate indicated that Bmp signalling played a crucial role in improving cell reprogramming in the pAAm gel condition at the early stages.

Further investigation of the protein-protein interaction network of DEGs at day 3 showed that a relatively unknown protein, Phactr3, was upregulated in the gel condition, but was not connected with other genes (**Figure S9b**, Supporting Information), reflecting that very little is known about its involvement in biological processes, especially in cell reprogramming. Phactr3 belongs to a novel protein family containing four Protein Phosphatase 1 and Actin Regulatory (Phactr) proteins (Phactr1-4) [43]. Besides the MRTF (myocardin-related transcription factor) family, these are the only proteins containing a highly conserved RPEL domain which binds with G-actin, which acts as an actin monomer sensor in cells and modulates cell migration. [44–48] Phactr proteins also bind with Protein Phosphatases (PPs), which regulate metabolism, cell cycle, and muscle contractility.[49–51] Interestingly, Phactr3 is the only known Phactr associated with nuclear non- chromatin structure[48], therefore having the potential to influence the reprogramming process critically, as nuclear non-chromatin structures plays a core regulatory role in gene expression.[52] We thus next sought to validate the role of Phactr3 under standard reprogramming conditions, believing that it may connect the earliest changes in cytoskeleton and metabolism, influencing the reprogramming process in the hydrogel condition.

qPCR analysis on MEF samples (inoculated on TCPS and 102 kPa for overnight) showed that Phactr3 expression was almost 2-times higher in the hydrogel condition compared to TPCS (**Figure 7a**). Phactr3 expression in MEFs from different embryos showed high variability but its upregulation in the pAAm condition was consistent for MEFs from each embryo. Immunostaining of Phactr3 also confirmed qPCR data (**Figure S11a**, supplementary information). These data confirmed that Phactr3 upregulation is an early event and the hydrogel *itself* can induce its expression without dox-inducible TFs. Interestingly, among the previously reported reprogramming facilitating factors (Bmp2, Fgf2 and Gata2)[37–39], only Bmp2 expression was significantly higher in MEFs incubated overnight on the pAAm substrate. Application of siRNA against Phactr3 for four days during the reprogramming culture efficiently reduced its expression in the cells **(Figure 7b**).

**Figure 7.**
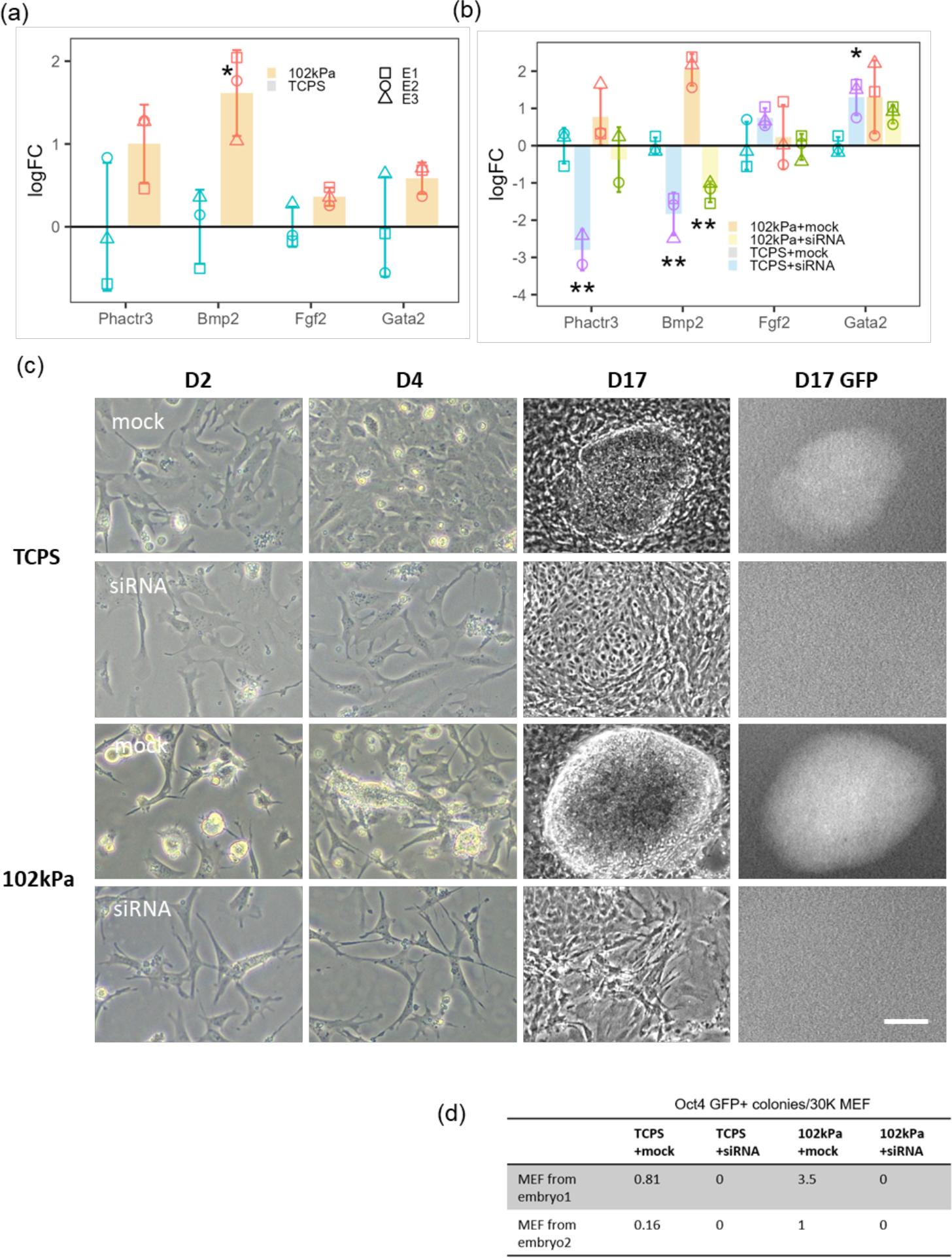
Regulation of cell reprogramming by Phactr3. (**a**) Phactr3 and other related gene expression (Log fold change (LogFC) relative to TCPS) in MEF inoculated in TCPS and 102 kPa overnight. (**b**) Same genes expression in MEFs reprogrammed for four days with scrambled pool of siRNA (mock) or Phactr3 siRNA in TCPS and 102 kPa. E1, E2 and E3 represents MEFs from different embryos and missing data point for any embryo represent not detected expression in qPCR. Line at y = 0 in (a) and (b) represents mean expression level in TCPS. Morphological observations in MEF reprogramming cultures at different time points (**c**) and Oct4 GFP+ miPSC colony counts at day 17 (**d**) in TCPS and 102 kPa with or without Phactr3 knockdown for four days. Data is represented as mean ± SD (n = 3) and “***”, “**” and “*” represents *p* < 0.001, *p* < 0.01 and *p* < 0.05, respectively. Comparisons are: TCPS vs pAAm for (a), and TCPS+mock vs TCPS + siRNA and 102 kPa + mock vs 102 kPa + siRNA for (b). Scale bar represents 250 µm (for D2 and D4) or 100µm (for D17 and D17 GFP). All images are in grayscale. miPSC colony counts are as # per 4.8 cm^2^.

For one MEF line, which showed the lowest Phacrt3 expression compared to other MEF lines, no expression of Phactr3 was detected under the siRNA condition. Upon Phactr3 knockdown with siRNA during a reprogramming culture for four days, Bmp2 expression was significantly downregulated **(Figure 7b**). Under Phactr3-knockdown reprogramming conditions, cells remained fibroblastic and failed to morph into epithelial-like cells and form clusters, a well-known characteristic change associated with MET at the early stage of reprogramming (**Figure 7c**).

Finally, suppression of Phactr3 for the first four days of reprogramming completely abolished the formation Oct4 GFP+ colonies in both TCPS and the 102 kPa gel condition (**Figure 7d**), indicating Phactr3 pays a crucial role in the early stages of reprogramming. These results confirmed that Phactr3 upregulation was one of the earliest events triggered by the hydrogel condition, and this upregulation appeared to drive Bmp2 upregulation, which ultimately lead to more efficient MET in the hydrogel condition, resulting in increased iPSC colony formation compared to TCPS.

## 3. Discussion

In this study, we reported that the pAAm gel had a pronounced effect on both the early and later stage of reprogramming (Fig. 1 and 2), whereas other studies thus far have only reported biomaterial-based modulation of the early stage.[17–20]

Downing and co-workers reported improvement of miPSC or hiPSC colony production by 4-times and 2-times, respectively, on 10 µm micro groove patterned PDMS surface compared to flat PDMS surface.[18] Choi and co-workers[17] transduced mouse fibroblasts with reprogramming transcription factors at day 1 on TCPS, plated them at day 2 on the hydrogels of different stiffness (0.1 kPa to 20 kPa) for one day and then replated the cells back onto TCPS for the remainder of the reprogramming period (another 18 days). They reported two times improvement in the reprogramming prone population (i.e., Thy1.2-/SSEA1+) and 2-times more miPSC colony production after the one-day of exposure of cells to the 0.1 kPa condition, compared to other stiffnesses. Caiazzo and co-workers[19] reported an improvement in reprogramming with very soft (0.6 kPa) 3D PEG-based hydrogels, that achieved 2.5-fold (at day 16) and 2-fold (at 6-weeks) more miPSCs and hiPSC colonies, respectively.[19] Kim and co-workers[20] only reported improvement in mouse iPSC production using methacrylated hyaluronic acid hydrogel with stiffness 0.1 kPa. These studies highlighted the substrate-mediated improvement of MET during the early stage of reprogramming for the overall improvement of the reprogramming outcome i.e., number of iPSC colonies.

In contrast to previous studies, we performed reprogramming continuously on 2D hydrogel substrates without the need for replating and observed two times more Thy1.2-/SSEA1+ murine cells at day 3 and four times more Oct4 GFP+ miPSC colonies at the end of reprogramming (Fig. 1). In the case of human fibroblasts reprogramming, we observed a pronounced impact in the later stages of reprogramming compared to the early stages under the hydrogel condition, resulting in 2- fold more hiPSC colonies and a 10-times greater yield of hiPSCs compared to TCPS in under 3 weeks (Fig. 2). We also observed that both mouse and human iPSCs from the pAAm condition had lower expression of primed ESC-markers and exogenous TFs used for reprogramming (i.e., OKSM or Sendai vector expression) compared to the TCPS condition (Fig. 3). 3D cell culture systems or patterned micro grooved substrates used for reprogramming in previous studies[18–20] are still challenging to be adopted in laboratories for routine production of iPSCs, which itself is a sensitive process. Overall, our study demonstrated 2D hydrogel-based improvement of both early and late phases of reprogramming and established an optimised pAAm hydrogel substrate (that can be easily made with a very low cost) platform as a robust 2D surface for accelerated manufacturing of quality iPSCs.

Another notable observation is that the hydrogel determined to be optimum for reprogramming in this study is much stiffer than the hydrogels reported in previous studies (< 1 kPa). [17,19,20] In contrast to the results reported by Choi and co-workers (lower % of Ssea-1+ population in stiffness > 4 kPa than < 1 kPa), we observed that hydrogels of all stiffness (1 kPa to 1.3 MPa) produced more Ssea-1+ population than TCPS at day 3 (Fig. S4b). Notably, Choi and co- workers analysed for the percent-Ssea-1+ population after only one day of exposure of the already transduced mouse fibroblasts (on TCPS) on the hydrogel substates, whereas we analysed at day 3 post transduction on the same substrates. After one day of exposure to their hydrogels, Choi and co- workers replated the cells onto TCPS for the remaining reprogramming period. However, we managed to perform the reprogramming on the hydrogel from start to finish, representing a very different, simpler approach. Our results indicate that, if enough time is given, stiffer hydrogels (in the range of kPa to MPa) have an equal potential to form increased numbers of reprogramming- prone populations at the early stage of reprogramming. We observed that cell growth rate was generally lower on hydrogels, and moreover that hydrogels below ≤ 16 kPa could not recover growth rates at later stages of reprogramming (Fig. S4a). Choi and co-workers investigated pAAm hydrogels of stiffness 0.1 kPa to 20 kPa, which fall in the range we found to not be supportive of cell growth, potentially providing reasoning for the need to replate the cells on TCPS for the remainder of the reprogramming period in their experiments. In contrast, we were able to identify an optimum hydrogel of a higher stiffness (102 kPa) which improved both early and late stages of reprogramming, without impacting final cell growth outcomes. It is worth noting that in previous works investigating 3D hydrogels for reprogramming (of stiffnesses < 1 kPa), a high number of cells (1-2 M/mL) was required to optimise the reprogramming outcome[19, 20] as very soft hydrogels impart growth limitations at low to moderate cell densities.[19] Improvements in reprogramming in our pAAm hydrogels compared to reports on 3D hydrogels was found to be slightly higher on the same comparators, being 4-fold versus 2.5-fold more Oct4 GFP+ mIPSC colonies generated (hiPSC colonies were measured using different markers). The 2D pAAm hydrogel format offers significant advantages over 3D alternatives, in being a low-cost, easily-made and scalable platform in itself, but also for further optimisation with the inclusion of other biophysical cues (e.g., microtopography[53], light-induced patterned region[54]) and small molecules.[55]

In this study, we have identified a novel mechanosensitive molecule, Phactr3, which was upregulated in the pAAm gel condition at the earliest time point, without any influence from the exogenous TFs expression, and confirmed it to be crucial for Bmp2-mediated MET and reprogramming (**Fig. 7**). In addition to Phactr3 upregulation, various genes related to metabolism (Hdac11, Pygl, Ppp1ca and Ppp2ca) were upregulated in MEFs exposed to the pAAm gel overnight (**Fig. S11b**, Supporting Information). As Phactr proteins bind with Protein Phosphatases (PPs), which regulate metabolism, cell cycle, and muscle contractility, [49–51], it is highly plausible that Phactr3 is also associated with the observed significant metabolic changes in the hydrogel condition at the early stage of reprogramming. Understanding the exact role of Phactr3 (and other Phactrs) on metabolism during reprogramming requires further investigation of the early-stage reprogramming process at shorter time spans (i.e., less than 3 days), as metabolism changes rapidly in these stages.[36] Future studies exploring the impacts of topography, 2D topography-patterned hydrogel or 3D gel conditions on Phactr3 expression and its impact on reprogramming and other cellular process outcomes would also be useful.

The large RNA-Seq dataset generated in the study contains six time point samples (day 0 MEF, reprogramming intermediates from day 3 and day 12, and Oct4 GFP+ iPSCs at day 17) from two substrate conditions. In this study, we explored only some of the possible molecular targets at day 3, leaving potential opportunity to explore more (e.g., role of non-canonical Wnt signalling, as Wnt7a was highly upregulated and its association with Phactr3) and other time points (e.g., from day 6, as from this point acceleration of reprogramming started in the pAAm gel condition).

Though we observed large improvements in the later stage of reprogramming, it is not known as to whether the improvement is independent of the early stage of reprogramming. Further, the distinct impact of the early and late-stage improvements on iPSCs characteristics, which was different between the gel and TCPS conditions, is not known. Future studies guided by the large RNA-Seq dataset will pave the way to answering these questions and potentially lead to the establishment of a robust reprogramming culture substrate and methodology for producing quality iPSCs in a more cost-effective manner than current gold-standard practices for regenerative medicine applications.

By using an optimised hydrogel substrate, we confirmed the notion by Tapia et al.[3] that faster kinetics and efficient reprogramming impacts the quality or characteristics of iPSCs. Even after the bulk passaging (more than 5-times) of mouse or human iPSCs on the TCPS, characteristics of iPSC produced in the pAAm hydrogel condition was found to be different compared to the TCPS-produced iPSCs (Fig. 3), presumably indicating different epigenetic status. We also observed that both mouse and human iPSCs from the pAAm condition had less exogenous expressions of the TFs used for reprogramming (i.e., OKSM or Sendai vector expression) compared to the TCPS condition. Higher growth rates of cells at the later period of reprogramming (**Figure 1d and 2e**) in the pAAm gel condition might be a reason for the accelerated loss of transgene expression. These data suggest that besides the observed significant influences over the ‘production’ outcomes of the reprogramming process, the stiffness of a hydrogel may also have greater influence over genetic change/insults that occur over iPSC culture periods.[3, 56] In this study, we did not assess the quality of iPSCs in a broader scope (e.g., immunogenic, genomic abnormalities, or epigenetic status) or long term propagation of iPSCs on the respective surfaces (i.e., hydrogel or TCPS) they are produced from. Therefore, studies that further compare the quality of iPSCs produced from pAAm and TCPS from a broader perspective would be important to pursue, as iPSC quality strongly determines differentiation efficiency and quality of the differentiated cells.[56].

## 4. Conclusion

In this study, by screening a series of 2D polyacrylamide hydrogels of varying stiffness, we found an optimum pAAm hydrogel condition (of stiffness 102 kPa) which significantly boosted both mouse and human iPSC production from fibroblasts compared to gold-standard TCPS-based protocols. This methodology permitted continuous reprogramming of fibroblasts on the hydrogel substrate, without the need for intermediate cell replating, even using Sendai virus methods for hIPSC generation. Using both experimental and time-course RNA-Seq data sets, we demonstrated that the 2D hydrogel condition accelerated the reprogramming processes and modulated both early and late stages reprogramming, ultimately producing more iPSCs (10-fold) over the same time period (18 days) that had distinct characteristics compared to those produced using conventional TCPS-based culture practices. Furthermore, we identified a novel mechanosensitive protein, phactr3, which critically impacted early-stage reprogramming and subsequently the end outcome of the reprogramming in both hydrogel-based and standard TCPS culture methods. These results indicate that the optimised 2D hydrogel platform discovered herein has significant potential as a robust platform for producing quality iPSCs more effectively. Future studies, supported by the large reprogramming time-course RNA-Seq data set made available by this study, focusing on the detailed assessment (e.g., immunogenic and epigenetic) of hydrogel-based modulation of iPSC quality, the impact of hydrogel surface modifications (e.g., photo-induce pattern, topography) on reprogramming outcomes, and the effect of the identified novel molecular target phactr3, will likely produce further insight and maturation of hydrogel-based methodologies for cell reprogramming.

## Acknowledgments

This work was funded in parts by the CSIRO Office of the Chief Executive Science Leader Fellowship (J.C.-W.), the Australian Research Council Discovery Grants Scheme (DP190101969) and the University of Queensland Vice Chancellor’s Strategy Funding Scheme under UQ StemCARE. This work was performed in part at the Queensland node of the Australian National Fabrication Facility, a company established under the National Collaborative Research Infrastructure Strategy to provide nano and microfabrication facilities for Australia’s researchers. This project was also part funded by a NHMRC Project Grant (GNT1051117), ARC Future Fellowship (FT180100674) and Sylvia and Charles Viertel Senior Medical Research Fellowship funds awarded to J.M.P.

## 5. Materials and Methods

### 5.1. pAAm hydrogels fabrication

Coverslips were cleaned in a Harrick Plasma Cleaner (PDC-002-HP, 45W) at 600 microns for 10 min. They were then soaked in prepared 0.4% (v/v) 3-(Trimethoxysilyl)propyl methacrylate) (M6514, Sigma-Aldrich) in distilled H2O (dH2O, pH balanced to 3.5 using glacial acetic acid) for 1 hr, before being washed twice with dH2O and dried in an oven at 120 °C for 1 hr. To increase gel stiffness from 1 kPa to 247 kPa, 40% (m/v) acrylamide (AAm) (A3553, Sigma-Aldrich) and 2% (m/v) bis-acrylamide (bis-AAm) (M7279, Sigma-Aldrich) stock solutions were mixed with dH2O to achieve the desired % (m/v) AAm and corresponding % (m/v) bis-AAm by maintaining their ratio at 29 : 1. For pAAm gel of 1.3 MPa, 180% AAm and 5% Bis-AAm solutions were mixed to achieve 80% (m/v) AAm and corresponding % (m/v) Bis-AAm. To form gel, 4 µl Tetramethylethylenediamine (T9281, Sigma-Aldrich) and 6 µl 10% (m/v) Ammonium Persulfate (A3678, Sigma-Aldrich) in dH2O were also added to AAm and Bis-AAm solution. Then, 150 µl drops of the complete solution were distributed on a hydrophobic glass slide (hydrophobized by using Sigmacote (SL2, Sigma-Aldrich)). The activated coverslips were then overlaid onto the drops and polymerization was continued for 20 – 30min. Coverslips with the gels were gently dislodged and washed twice with PBS and stored at 4 °C.

### 5.2. Rheology of the gels

For oscillatory rheology, gels were formed over non-activated coverslips by distributing 300 µl of gel solution. Gels were peeled off from the coverslips and placed between the plates (top plate diameter 8 mm) of an oscillatory rheometer (Discovery HR-2, TA instruments). Gel excess was scraped off using a scalpel and the top plate was lowered until a small bulge around the gel was observed. 3 µl of PBS was distributed around the gel before starting each rheological measurement to keep the gel hydrated. For rheological measurements, strain sweeps using strain range (0.01% or 10%) at 10 rad/s frequency were performed to obtain the Storage (*G’*) and loss (*G’’*) modulus vs strain curve. *G’* and *G’’* were then determined at various frequencies (range 1 rad/s to 100 rad/s) by using the specific strain value obtained from the linear viscoelastic range of the curve. Finally, G’ and *G’’* values at 10 rad/s were reported as the gel’s rheological properties. From these values, the Young’s modulus (*E*) was calculated using 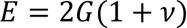, where 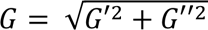 and *ν* is Poisson’s ratio 0.457.[57]

### 5.3. ECM immobilization using Sulfo-SANPAH and L-DOPA

ECM immobilization with Sulfo-SANPAH treatment was performed according to a published protocol.[58] In brief, 50 mg Sulfo-SANPAH (sulfosuccinimidyl 6-(4’-azido-2’- nitrophenylamino)hexanoate) (cat. 22589, ThermoFisher Scientific) was dissolved in 1 ml DMSO and diluted 25-times in Milli-Q water, resulting in a 2 mg/ml Sulfo-SANPAH solution. 300 µl of this solution was added to the gel surface and placed under a UV light in a biological safety cabinet for 30 min. Gels were then washed twice with PBS and incubated with the desired ECM (0.2% Gelatin in PBS, G1890 Sigma-Aldrich) overnight at 4 °C. Gels were prepared for cell culture by washing once in PBS, rinsing in 70% ethanol for 2 min, washing in PBS twice, UV-sterilised for 30 min and incubating with culture medium for 1 hr. For ECM conjugation using L-DOPA (3, 4- Dihydroxy-L-phenylalanine, D9628, SIGMA),[23] 2 mg ml^-1^ L-DOPA solution was prepared using 10 mM Tris-HCL buffer (pH balanced to 10.2 using 1 M NaOH) by mixing gently on a roller shaker for 30 min under no-light conditions. The solution was filtered to remove undissolved L- DOPA, distributed over the gels, and then incubated for 30 min under no-light conditions. Gels were then conjugated with ECM and prepared for cell culture as per the procedure described for Sulfo-SANPAH above.

### 5.4. MEF reprogramming

Murine Embryonic fibroblast (MEF) cultures containing doxycycline inducible O (Oct4) K (Klf4) S (Sox2) M (Myc) cassette and endogenous Oct4 driven GFP expression locus were provided by Dr Jose M. Polo group at Monash University.[24] MEFs were expanded for one passage (P1) in MEF medium (High-glucose DMEM, 10% FBS, sodium pyruvate, L-Glutamine, penicillin-streptomycin, non-essential amino acids, b-mercaptoethanol) for preparing cell stocks in FBS with 10% DMSO. These MEFs (inoculated at 25k cm^-2^) were expanded for 3 days maintaining < 90% confluency before inoculating them for reprogramming at 2000 cells cm^-2^ on the gel or TCPS conditions in ESC medium (KODMEM with 15% FBS, L-Glutamine, penicillin-streptomycin, non-essential amino acids, b-mercaptoethanol and 1000 U ml^-1^ LIF) and 2 ug ml^-1^ doxycycline (cat. 33429 Sigma-Aldrich). MEFs were reprogrammed in the presence of doxycycline for 12 days and without doxycycline for the last 5 days. At the end the reprogramming, Oct4 GFP+ iPSC colonies were counted manually under fluorescence microscope and Oct4 GFP+ cells were quantified by flow cytometry.

### 5.5. Human fibroblast reprogramming

To validate the efficacy of 102 kPa pAAm gel to improve human cell reprogramming, we utilised a modified Sendai-based transfection protocol to reprogram human dermal neonatal foreskin fibroblasts (hDFn). In the protocol, hDFns were inoculated at low density (4k cm^-2^) on the substrate, incubated overnight before transfecting with Sendai, and the reprogramming culture was continued without replating for 18 days. We confirmed that transfection of cells at low density with Sendai did not affect cell viability (Figure S5b, Supporting Information). Two different lots (lot 1355434 and 1767629) of human dermal neonatal foreskin fibroblasts (hDFn) were purchased from Thermo Fisher Scientific (cat. C0045C) and expanded twice according to manufacturer instructions. These hDFns (inoculated at 35k cm^-2^) were grown for 3 days (maintaining < 90% confluency) in MEF medium before inoculating them for reprogramming at 4000 cells cm^-2^ on the gels or TCPS. For human cell reprogramming, gels or TCPS were coated with vitronectin (0.2 µg cm^-2^, A14700, Thermo Fisher Scientific) instead of Gelatin. After overnight incubation, cells were transduced with Sendai virus (CytoTune – iPS 2.0 Sendai Reprogramming Kit, A16517, Thermo Fisher Scientific) using KOS multiplicity of infection (MOI): Myc MOI : Klf4 MOI = 5:5:6 in MEF medium. After 24 hrs (Day 1), culture medium was exchanged with E8 medium (cat. A1517001, Thermo Fisher Scientific). Culture medium was exchanged again on Day 2 and every other day after that. From Day 2 to Day 6, reprogramming was continued in E8 medium and from Day 6 to Day 18 in E7 medium (E6 medium, cat. A1516401, Thermo Fisher Scientific + 100ng ml^-1^ bFGF, cat. 100-18B, Peprotech). On Day 18, the reprogrammed culture was stained for Alkaline phosphatase (AP) activity. AP+ hiPSC colonies were counted manually and SSEA4+/TRA1-60+ hiPSCs were quantified by flow cytometry.

### 5.6. Alkaline phosphatase (AP) staining

Cells were fixed with 4%PFA solution for 15min and washed once with PBS. Afterwards, a filtered solution containing 0.2mg/ml Napthol AS-MX Phosphate disodium salt (Sigma-Aldrich) and 1mg/ml Fast Red TR Salt hemi (Zinc Chloride) salt in 100mM Tris-HCl buffer (pH 9.2) was distributed over the cells and incubated for 20min. Finally, cells were washed twice with PBS.

### 5.7. Flow cytometry

Mouse cells and human cells were dissociated with 0.25% Trypsin-EDTA (cat. 25200056, Thermo Fisher Scientific) and TrypLE (cat. 12604013, Thermo Fisher Scientific), respectively. Dissociated cells were washed once with 1% BSA in PBS before incubating with primary antibodies for 30 min. The following primary antibodies were used for mouse and human cells: Anti-Thy-1.2 PE-Cyanine (1:400, cat. 25-0902, eBioscience), Anti-SSEA-1 Biotin (1:400, cat. 13-8813, eBioscience), Anti- CD13 PE-Cyanine (2.5 µl/test, cat. 561599, BD Pharmingen), Anti-SSEA-4 PE (2.5 µl/test, cat. 330406, Biolegend) and Anti-TRA-1-60 Alexa Fluor 647 (2.5 µl/test, cat. 330606, Biolegend). Cells were washed twice with 1% BSA in PBS and incubated with the secondary antibody Streptavidin APC (1:200, cat. 17-4317, eBioscience) for 30 min if necessary. Finally, cells were washed twice in 1%BSA in PBS containing propidium iodide and passed through a 40 µm cell strainer to achieve single cell suspension. Cells were analysed in a LSRII (BD Bioscience) and sorted by Influx cell sorter instrument (BD Biosciences).

### 5.8. RNA analysis

Total RNA isolation from cell pellets was performed according to manufacturer’s instructions for RNeasy Mini or Micro kit (cat. 74104 or 74004, Qiagen). RNA was eluted from the columns using 20 µl of RNase-free water and quantified using Nanodrop.

### 5.9. RNA-Seq

Total RNA samples (RIN > 8) were sequenced on HiSeq 1500 (Illumina Inc.) using HiSeqV2 Rapid chemistry according manufacturer’s instructions in Monash Health Translation Precinct RNA-Seq facility. All RNA-Seq data will be available from the GEO repository.

### 5.10. Bioinformatics analysis

#### 5.10.1. RNA-Seq raw data processing

Reads were aligned using the custom genome and gtf file (gencode vM4) provided by Dr Jose Polo[24] with the STAR aligner in per-sample 2-pass mapping mode (version 2.5.1b).[59] The number of reads maps to each gene was counted using htseq-count248 (version 0.6.1) with python version 2.7.8. Since our RNA-seq data is from a strand specific assay, the stranded parameter was set to reverse. The order parameter was also set to “pos” since the alignment was sorted by alignment position. Count data was converted to RPKM values using the function “rpkm” from edgeR (version 3.18.1 and limma version 3.32.5) with norm.lib.sizes set to TRUE (default). The lengths of each gene were acquired using featureCounts on one sample and the custom GTF file. The RPKM values were log2 transformed with one added to each value to prevent NAs.

#### 5.10.2 Enrichment analysis of MIKKELSEN gene sets in GSEA

To perform enrichment of various MIKKELSEN gene sets in GSEA, which contain differentially regulated or methylated genes in MEF, mESC, fully and partially reprogrammed cells,[26] C2 curated gene sets for mouse were downloaded from http://bioinf.wehi.edu.au/software/MSigDB/, and a camera test was performed in R using the gene sets and DEGs between Oct4 GPF+ cells in TCPS and pAAm. Though iPSC (passage 0, i.e., P0) generated from TCPS and pAAm gel clustered quite closely, over 3000 genes (FDR = 0.05) were differentially expressed between them at Day17. Although, MIKKELSEN_PLURIPOTENT_STATE_UP/DOWN (genes up/down regulated in the iPSC, and ESC compared to the parental lineage-committed and partially reprogrammed cell lines) regulation was not different (FDR > 0.05), MIKKELSEN_DEDIFFERENTIATED_STATE_UP and DOWN (genes up/down in partially reprogrammed, iPSC and ESC compared to parental lineage-committed cell lines) were up and down regulated in iPSCs produced on pAAm gels compared to TCPS (Figure S6a, Supporting Information). Along with the downregulation of gene sets which are inactive in ESC and iPSC (marked by H3K27ME3 and both H3K4ME3 and H3K27ME3), iPSC in the pAAm condition downregulated gene sets active in MEF (MIKKELSEN_MEF_LCP_WITH_H3K4ME3) compared to TCPS (Figure S6a, Supporting Information).

#### 5.10.3. KEGG pathway analysis

KEGG pathway analysis was performed using kegga function in limma (v.3.28.14) available in R. FDR < 0.05 and *p* < 0.05 were used to filter DEGs and differentially regulated pathways, respectively.

#### 5.10.4 Principal Component Analysis (PCA)

PCA plot was generated using plotPCA in DESeq2 in R (version 3.4.1).

#### 5.10.5 Identification of molecules accelerating reprogramming in pAAm gel

First, we performed STRING protein-protein interaction network analysis of the DEGs (FDR < 0.05, logFC > 1.5) at D3 by classifying them under various GO terms: focal adhesion, cytoskeleton organization, cellular response to hypoxia, tight junction, TGF-beta signaling pathway, stem cell population maintenance, epithelium development, nucleus and metabolic process (Figure S8,

Supporting Informaiton) using Cytoscape (v3.6.1). These GO terms were relevant to the essential up/down regulated KEGG pathways at D3 identified by GAGE analysis.

It is well known that when cells are placed on a substrate, they probe the surface by forming and dislodging focal adhesions and adapt with the mechanical environment by changing cytoskeleton organization and extracellular matrix.[60–65] Indeed, cells on pAAm undergo large changes in cytoskeleton organization indicated by downregulation of various actin binding proteins[66] e.g., Actn1, Cap2 and Tpm2 (**Figure S9, S10**, Supporting Information).

Cells on pAAm express higher Fgf2 compared to TCPS, which was strongly associated with lower Fn1 and collagen expression (except Collagen15 and Collagen6) (**Figure S9, S10**, Supporting Information). Interestingly, Collagen15 and Collagen6 are the non-fibrillar collagen which are presents in epithelia basal lamina and knockdown of Collagen6 and associated upregulation of Fn1 caused loss of epithelia cell shape.[67, 68] Fgf2 has the potential to induce reorganization and disruption of actin cytoskeleton through PI3K, Rho and Cdc42 and reduce the Fn1 expression and Fn1-mediated collagen 1 family expression. [69–72] Jiao and co-workers showed that collagen downregulation in the presence of Fgf2 at the early stage of reprogramming improved cell reprogramming outcomes,[38] which presumably played a major role in improving cell reprogramming outcome in the pAAm condition in this work.

Interestingly, we observed that Gata2 expression was upregulated across the whole reprogramming period in pAAm compared to TCPS (**Figure S9, S10**, Supporting Information). Arhgap proteins (Rho GTPase Activating Protein) activates the RhoA pathway to modulate cytoskeletal organisation.[73] Various Arhgap proteins (Arhgap22 and Arhgap23) were downregulated in the pAAm condition on day3, and like Arhgap35, downregulation of these might have induced upregulation and trans localization of Gata2 for cells on pAAm.[73, 74] Recently, the Gata transcription factor family has been shown to be similarly effective as or better than Oct4 to induce reprogramming. Notably, among Gata TFs, Gata2 showed more effectiveness to improve reprogramming.[39] Therefore, besides Fgf and Bmp signalling pathways, Gata2 presumably played an important role in the reprogramming outcomes observed in the pAAm gel condition.

### 5.11. Knockdown of Phactr3 with siRNA

siRNA against Phactr3 or mock scramble siRNA at 4nM was prepared in Opti-MEM® I Reduced Serum Medium (Life Technologies 51985-026) with 16ul Lipofectamine® RNAiMAX Transfection Reagent (Life Technologies 13778075)/ml solution and incubating the solution at room temperature for 20mins. Then, 1ml of this solution was mixed with 3ml MEF culture medium and on day 1 of routine culture of MEF, at 40-50% confluency in T25 flasks, cells were incubated with 4ml containing siRNA against Phactr3 (ON-TARGETplus Mouse Phactr3 (74189) siRNA – SMARTpool, Lot-046898-01-0005, Dhramacon) or mock scramble siRNA (DHA-D-001810-10-05 ON-TARGETplus Non-targeting Pool) at 2 nM and cultured for another two days before using them in reprogramming experiments. At the start of reprogramming experiments, MEFs were inoculated on TCPS or pAAm hydrogel in MEF media without siRNA. On the following day, media was changed to reprogramming media containing target siRNA or scramble siRNA. On day 2, cells were transfected with respective siRNAs for another two days and after that cells were reprogrammed as usual.

### 5.12. Immunostaining

Cells were rinsed twice in PBS, fixed in 4% PFA for 20mins, washed in PBS twice for 5min each wash, permeabilized with 0.2% Trition for 10mins, washed again twice in PBS, blocked with 5% goat serum in PBS for 30min, and incubated with Phactr3 antibody (PA5-65682, 2µg/ml) overnight. Cells were then washed thrice in PBS, incubated with secondary antibody (4 µg/ml) and/or Alexa Fluor 488 Phalloidin (A12379, 400 times diluted) for 90mins, washed thrice in PBS, stained with DAPI (2 µg/ml) for 5 min, and rinsed thrice in PBS. Cells were visualised within three days with normal fluorescence or confocal microscopy.

### 5.13. qPCR

Total RNA and cDNA were synthesized from the samples using RNeasy mini kit and QuantiTect Reverse Transcription Kit according to the manufacturer’s instructions (Qiagen). qPCR was performed in a Biorad Thermocycler with a reaction volume of 20 µl which constitutes primers (0.5 µM), cDNA (equivalent to 20 ng RNA) and PowerUp SYBR Green Master Mix. Cycling condition was 50 °C for 2 mins, 95 °C for 2mins, 95 °C for 15s and 60 °C for 1 min. The last two heating steps were repeated for 45 times. Primers are available upon request.

### 5.15. Statistical analysis

All values are expressed as the mean ± SEM if otherwise not stated. The data were analyzed using Student’s *t*-test or the ANOVA test. A *p* < 0.05 was considered statistically significant. GraphPad Prism (GraphPad Software Inc., San Diego, California, USA) was used for these analyses.

## Supplementary figures

**Figure S1.**
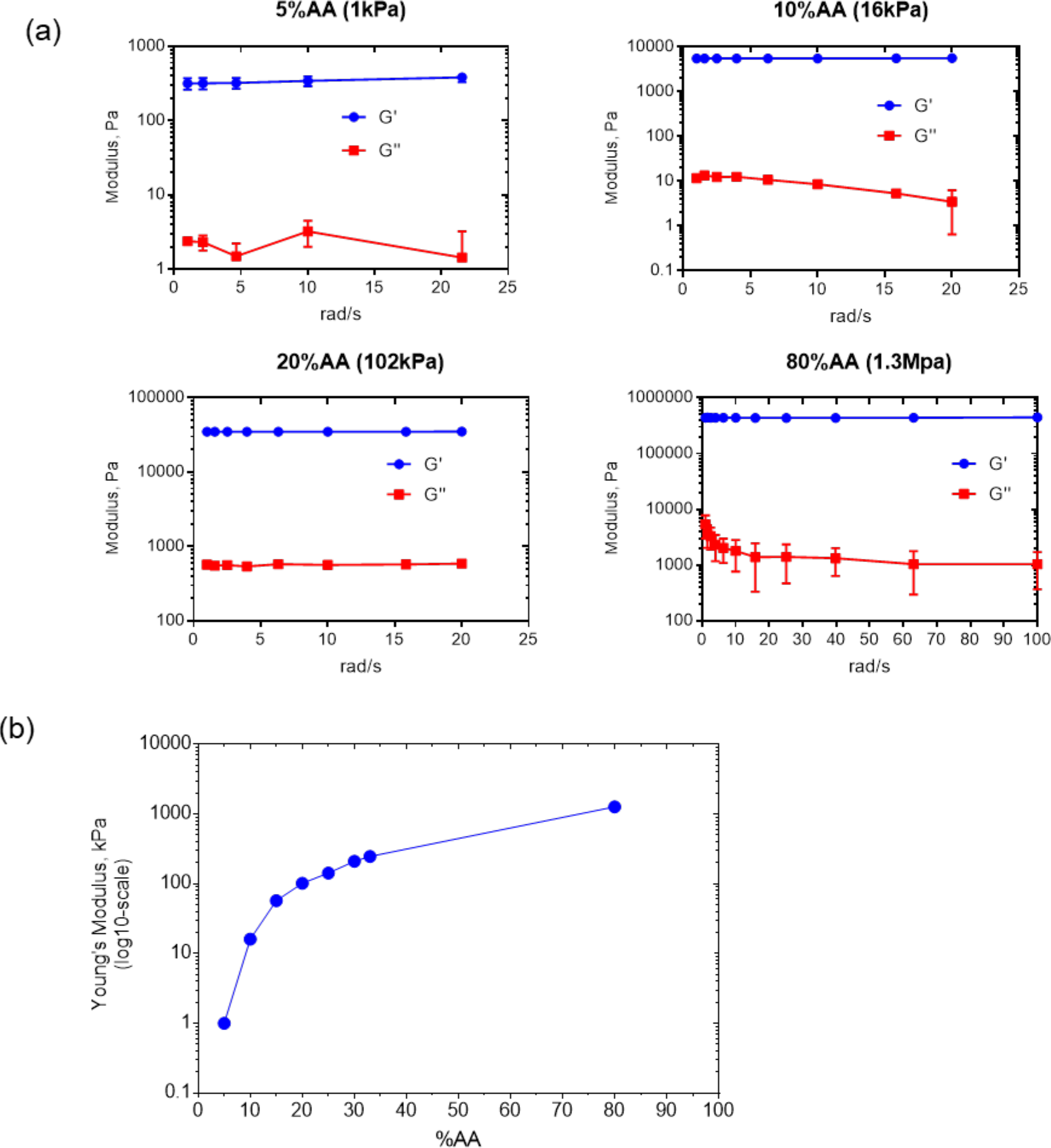
Rheological characterisation of polyacrylamide hydrogel (pAAm). (**a**) Storage (*G’*) and loss (*G’’*) modulus, measured by oscillatory rheometer, for representative pAAm gels prepared by varying acrylamide and bis-acrylamide amount in 29:1 ratio. Significantly lower value *G”* indicated mostly elastic behaviour of these gels. (**b**) Young’s Modulus (*E*) of pAAm gels calculated by using shear modulus (*G*) and Poisson’s ratio (ν = 0.457, as described in method section) (n = 2 or 3). Error bar represents Mean ± SD.

**Figure S2.**
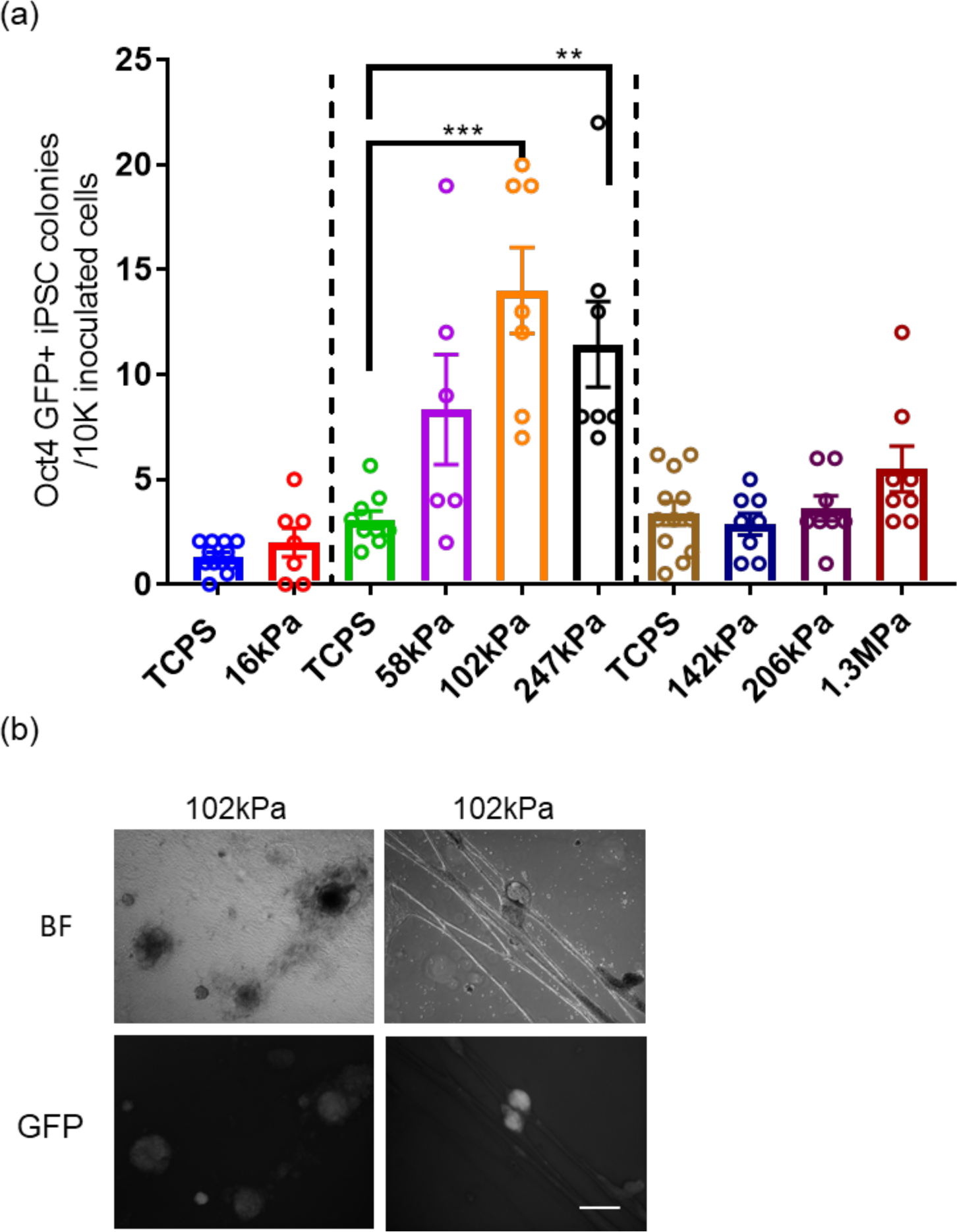
MEF reprogramming on pAAm gels of various stiffness (conjugated with Sulfo-SANPAH for ECM immobilization). (**a**) Number of Oct4 GFP+ miPSC colonies counted at day 17 on TCPS and pAAm gel of various stiffness. Data from different batches of experiment is presented together. In all cases, MEF from three different embryos and at least two technical replicates were used. Data is represented mean±SEM and “***” and “**” represents *p* < 0.001 and *p* < 0.01, respectively. (**b**) Bright field and corresponding GFP channel showing miPSC colonies adhered on the surface of the 102 kPa stiff hydrogel (left) and with the detached cell layer (right) over the same stiff hydrogel. Scale bar represents 500 µm.

**Figure S3.**
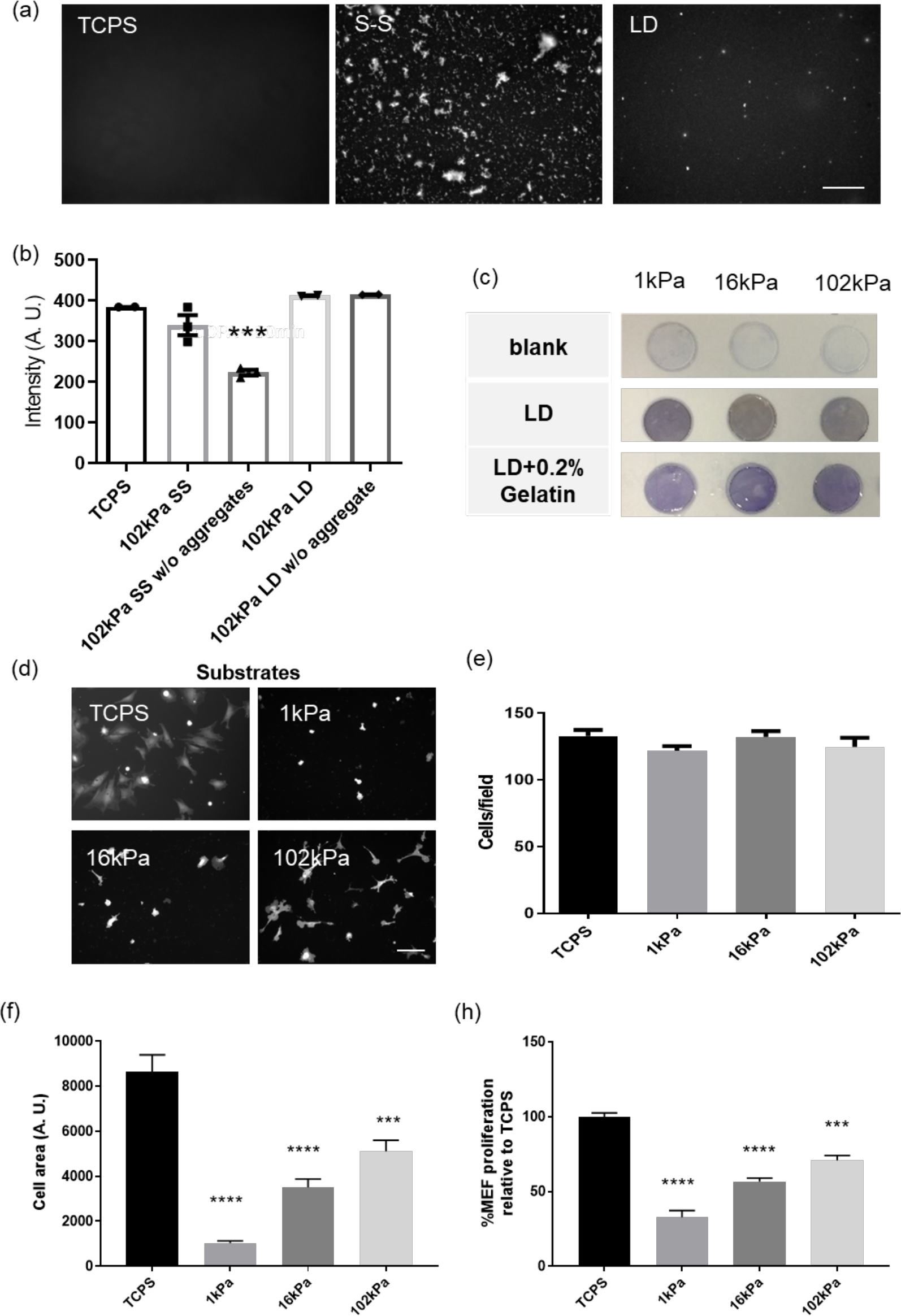
Characterisation of ECM immobilization on pAAm gel using Sulfo-SANPAH (SS) or L-DOPA (LD). (**a**) Images showing immunostaining of Col1A that was deposited on TCPS and pAAm (E = 102 kPa) treated with SS, LD (n = 2 or 3; scale bar: 125 µm). (**b**) Intensity of the images quantified by ImageJ (n = 2 or 3). Coomassie blue assay showing the amount of Gelatin deposition on hydrogel of various stiffness (**c**). Representative images showing MEF morphology on different substrates after overnight incubation (d; scale bar: 250 µm). Cells were stained with Calcein AM or Dil stain. Cell number per field (**e**) and cell area (**f**) after overnight incubation, and relative cell number quantified by presto blue assay at day3 (**g**). Data is represented as mean ± SEM and “****”, “***”, “**” and “*” represents *p* < 0.0001, *p* < 0.001, *p* < 0.01 and *p* < 0.05 compared to TCPS, respectively.

**Figure S4.**
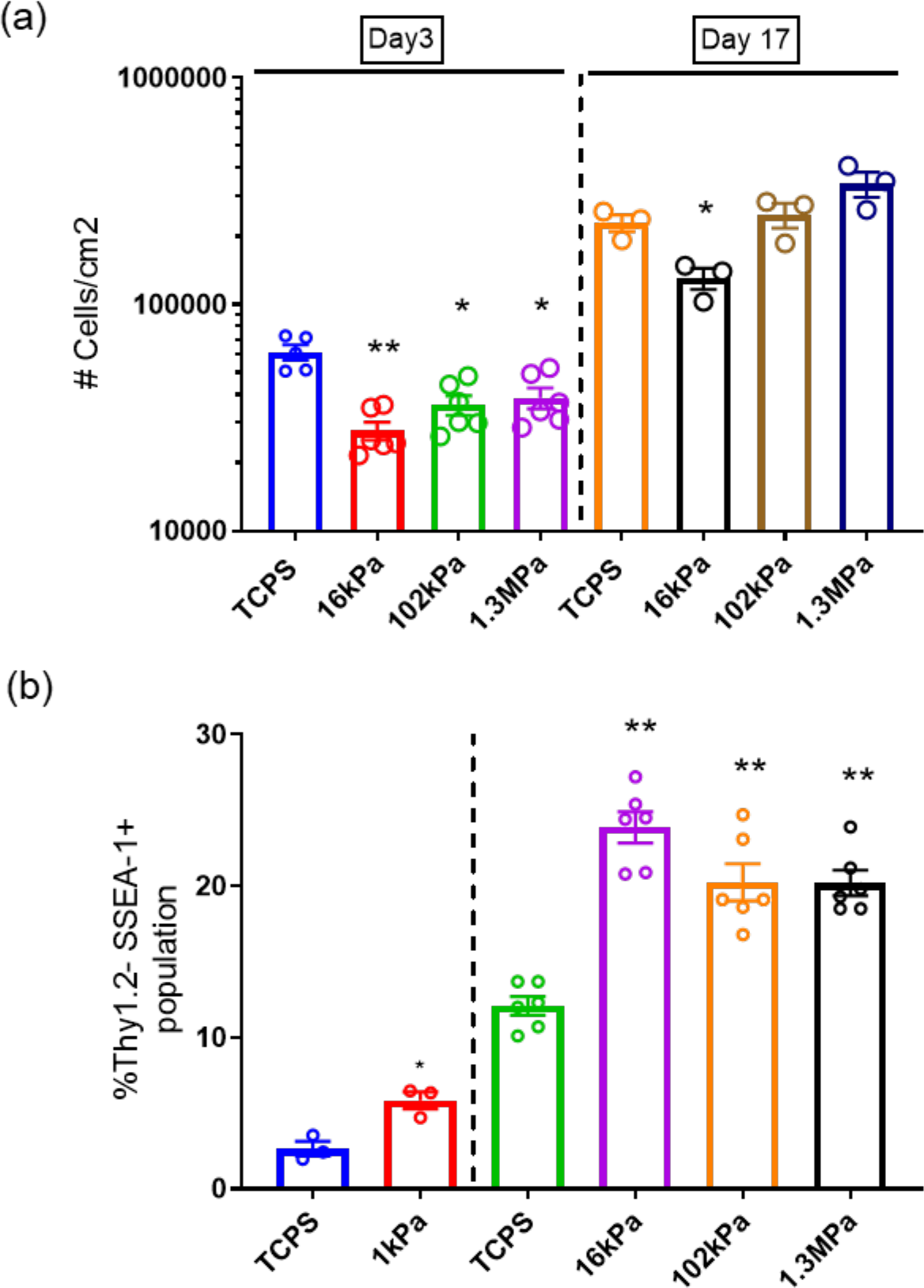
(**a**) Cell growth decrease (quantified by cell number) at early period (Day 3) of reprogramming but became similar to the TCPS at the end of reprogramming (Day 17) on pAAm gels for E ≥ 102 kPa. Soft substrate (*E* = 16 kPa) was not able to recover the initial drop-in growth rate at the end of reprogramming. (**b**) Percentage of cells undergoing reprogramming (Thy1.2- SSEA1+ cells) at Day 3 on TCPS and pAAm gel of various stiffness. Data from different batches of experiment is presented together. For all data, MEF from three different embryos and at least two technical replicates were used. Data is represented as mean ± SEM and “***”, “**” and “*” represents *p* < 0.001, *p* < 0.01 and *p* < 0.05 compared to TCPS, respectively.

**Figure S5.**
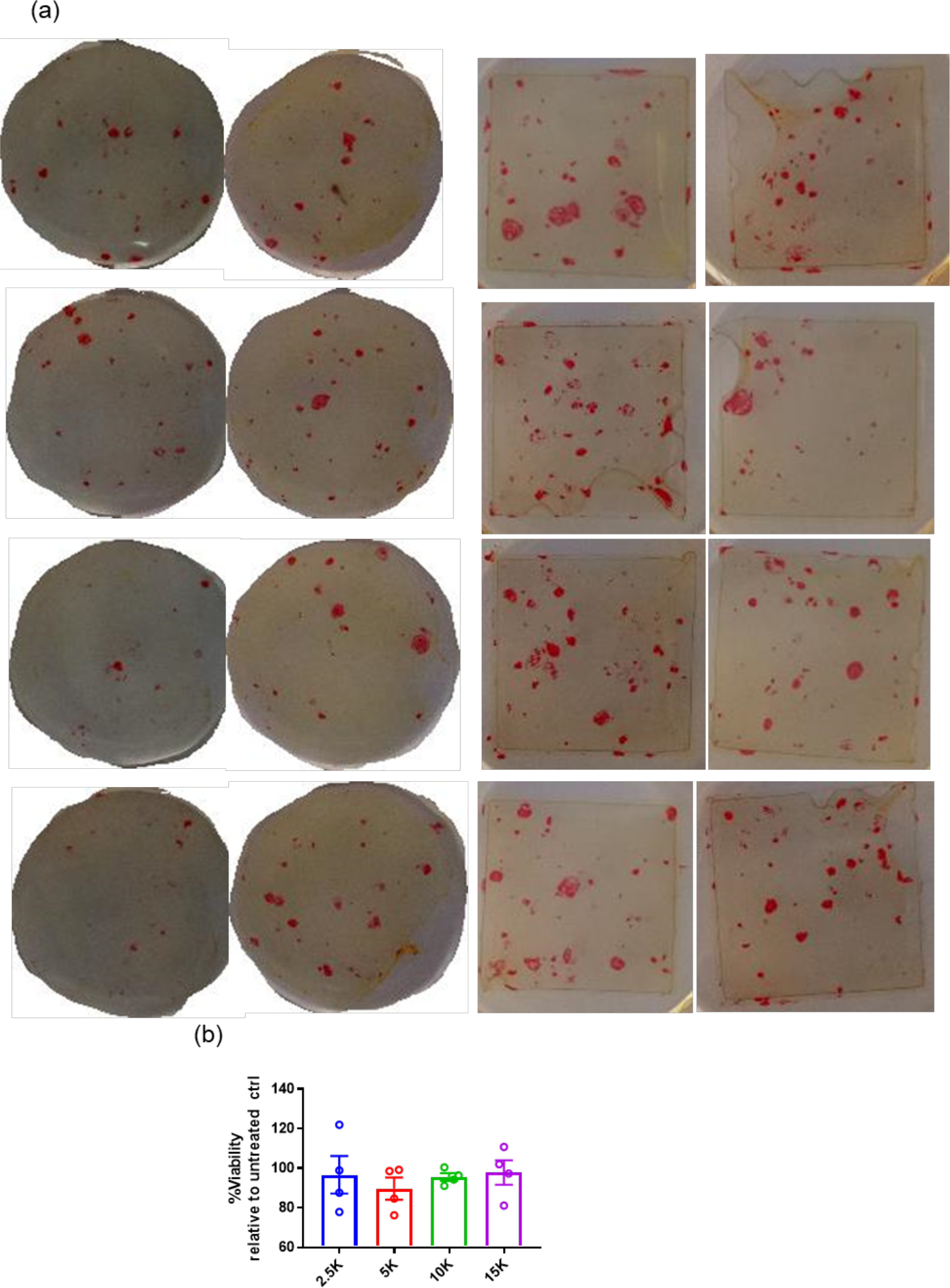
(a) Images of ALP+ (red) hiPSC colonies at day 18 in TCPS (first two columns) and 102 kPa (last two columns). Each well of TCPS represent a well of 12-well plate and for hydrogel each represent a 22 mm x 22 mm coverslip. **(b)** Viability of hDFn (inoculated at different densities) at day4 after inducing them with sendai virus for reprogramming. Data is represented as mean ± SEM.

**Figure S6.**
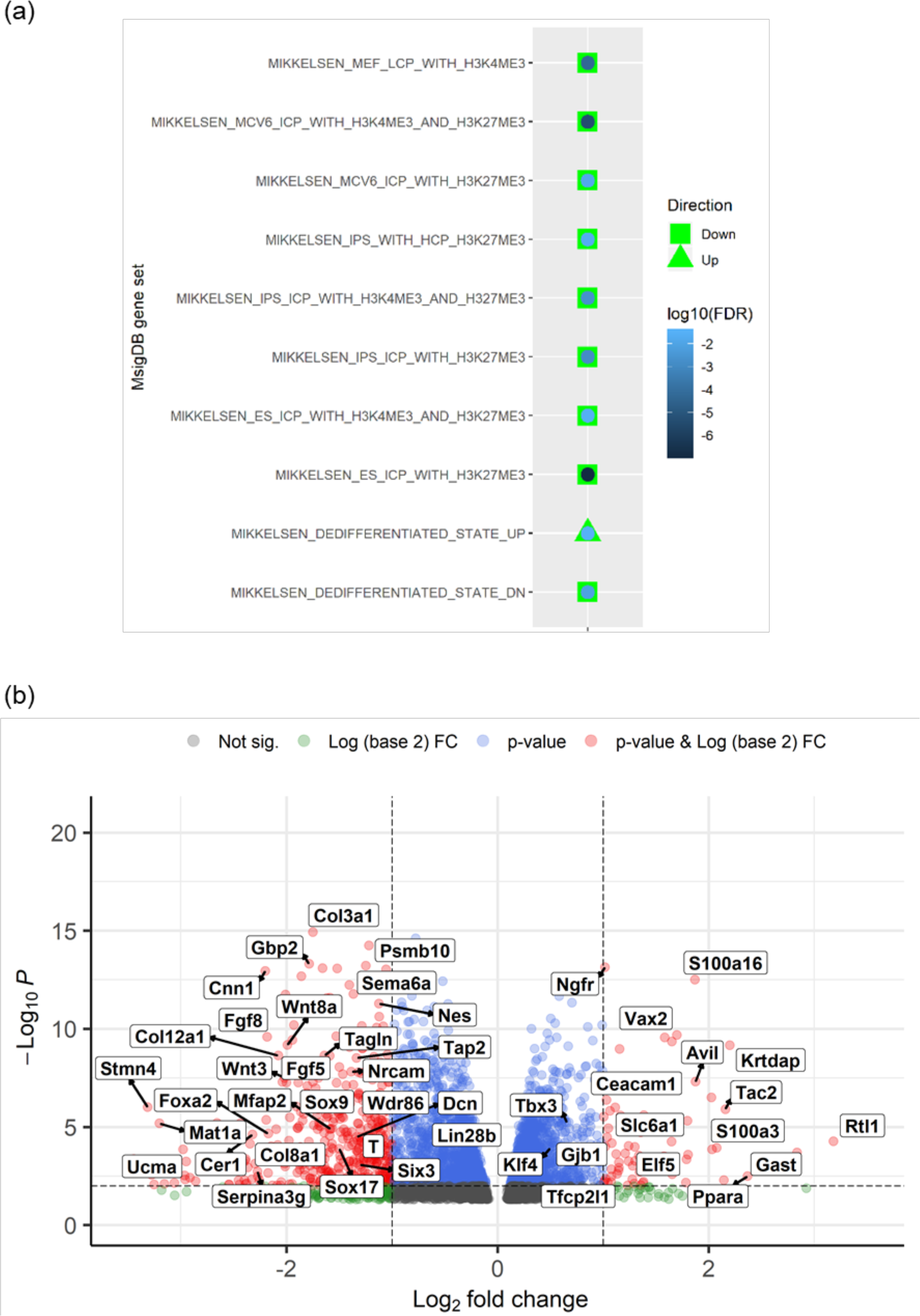
(a) Up or Down regulated Mikkelsen gene sets in MSigDB, which are related cell reprogramming to pluripotency. **(b)** Volcano plot of DEGs at day 17 between 102 kPa and TCPS (**b**). Horizontal and vertical dotted lines represent p cut-off value (0.01) logFC cut-off value (1), respectively.

**Figure S7.**
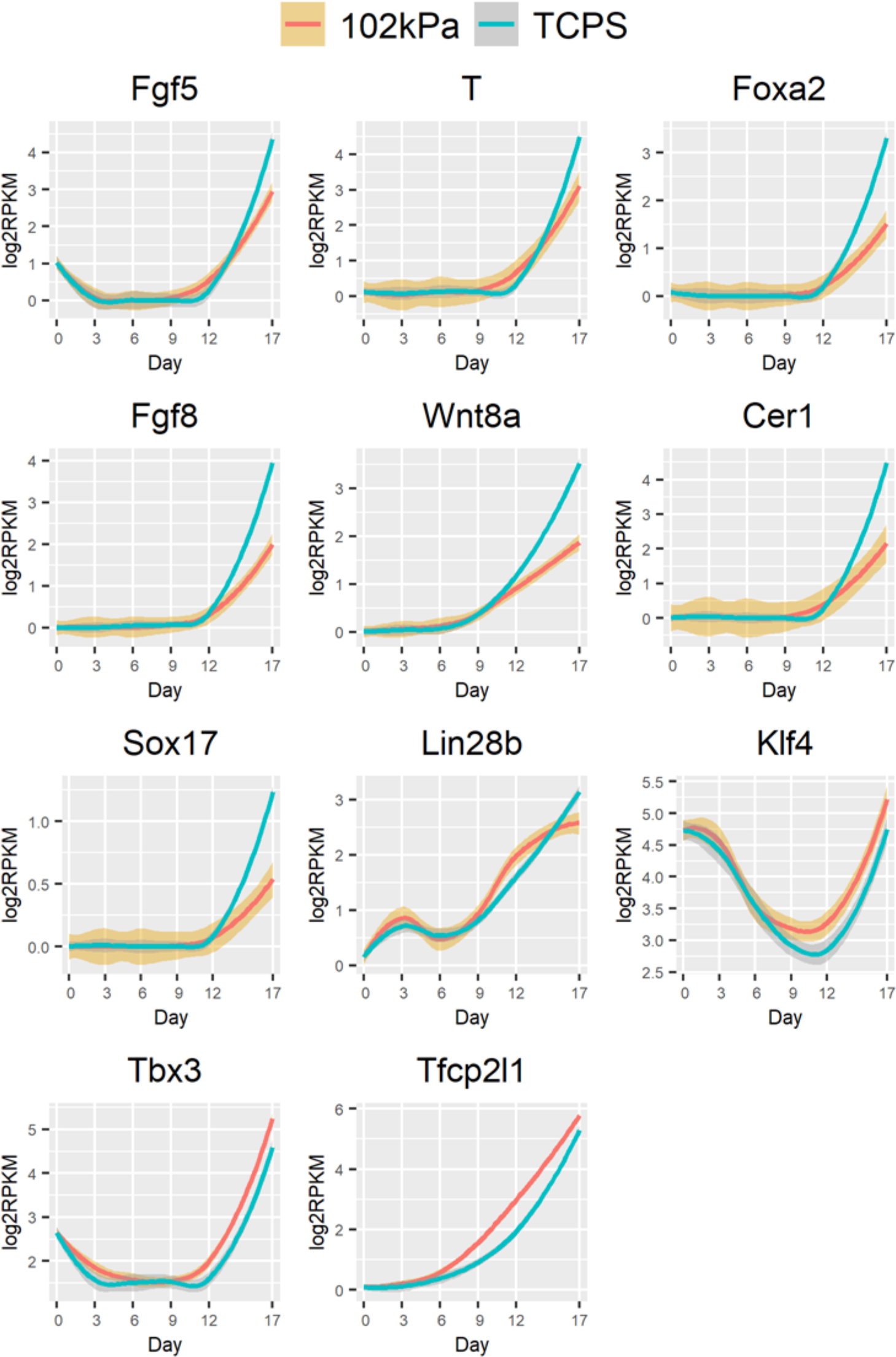
Expression profile of various genes that were mentioned in the main text. Band enclosing expression profile represent 95% confidence interval.

**Figure S8.**
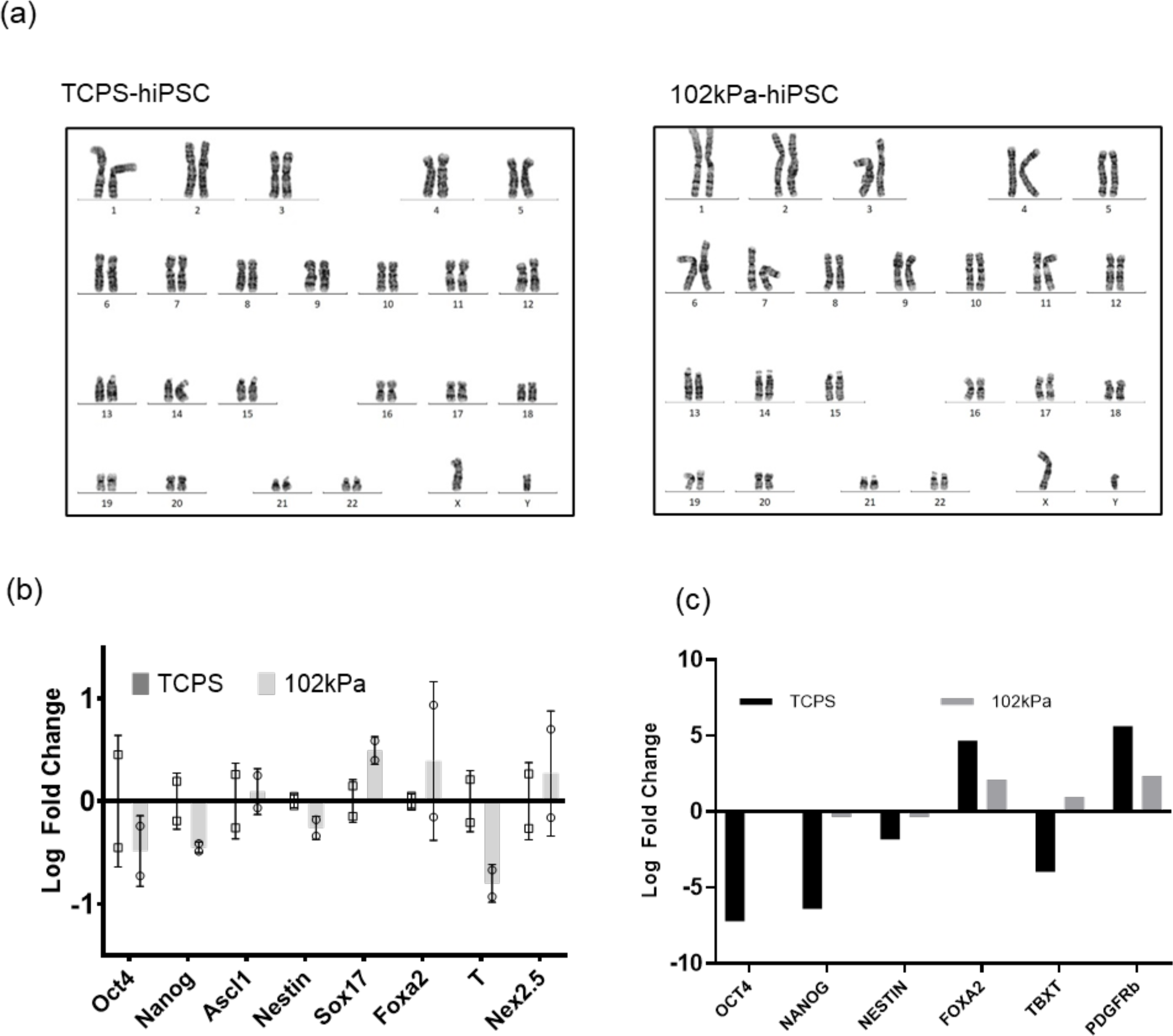
(**a**) Karyotype of hiPSC generated in TCPS and pAAm. (**b**) Expression level of PSC markers (Oct4/OCT4 and Nanog/NANOG) and differentiation makers (remaining ones) in **(b)** mEB (n = 2, relative to TCPS) and **(c)** hEB (n = 1, relative to expression in hEB formed with H9 hESCs). Line at y=0 represents mean expression in TCPS. mEBs or hEBs were formed from miPSC or hiPSC derived in TCPS and pAAm and differentiated for 8-days in 20% Knockout Serum containing medium. Data is represented as mean ± SD.

**Figure S9.**
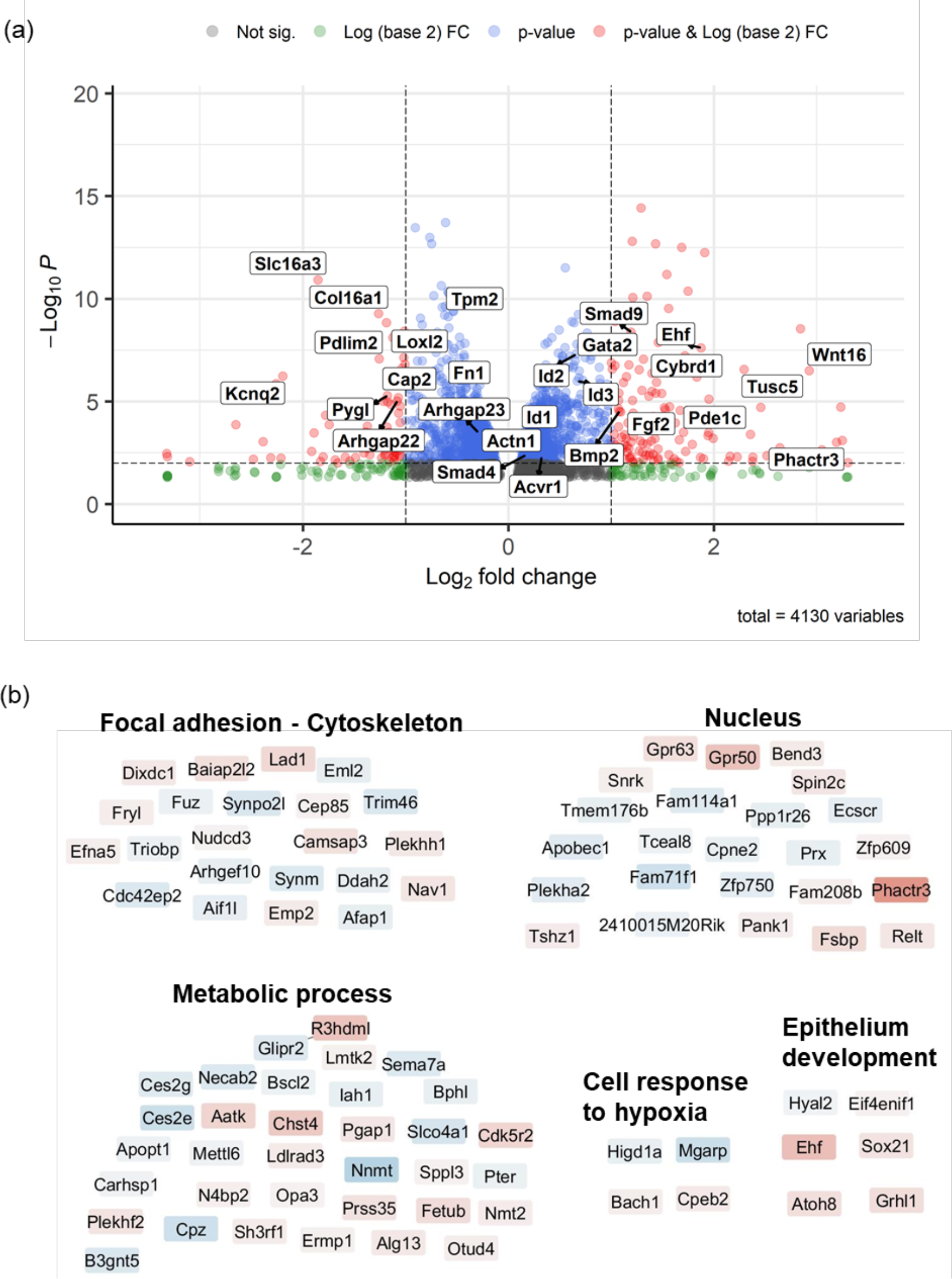
(**a**) Volcano plot of DEGs at day 3 between 102 kPa and TCPS. Horizontal and vertical dotted lines represent *p* cut-off value (0.01) logFC cut-off value (1), respectively. (**b**) Network view of lowly connected (relationship retrieved from STRING database) DEGs (between pAAm and TCPS at day3) in Cytoscape. DEGs (FDR = 0.01) were grouped together according to their biological function or location in a cell as described by the corresponding GO terms (bold text). Colour indicates relative log-fold change, red is higher, and blue is lower.

**Figure S10.**
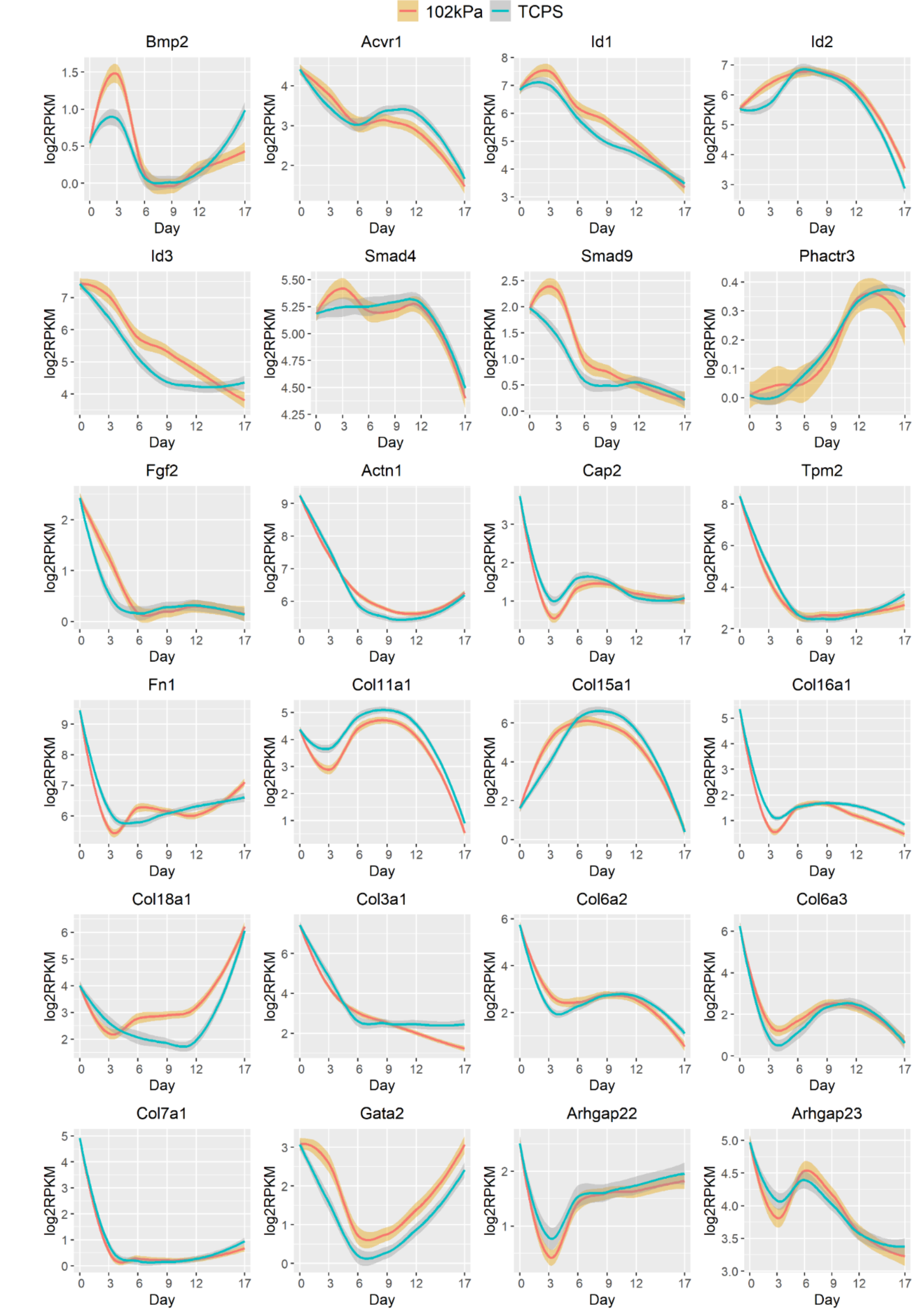
Expression profile of various genes that were mentioned in the main text. Band enclosing expression profile represent 95% confidence interval.

**Figure S11.**
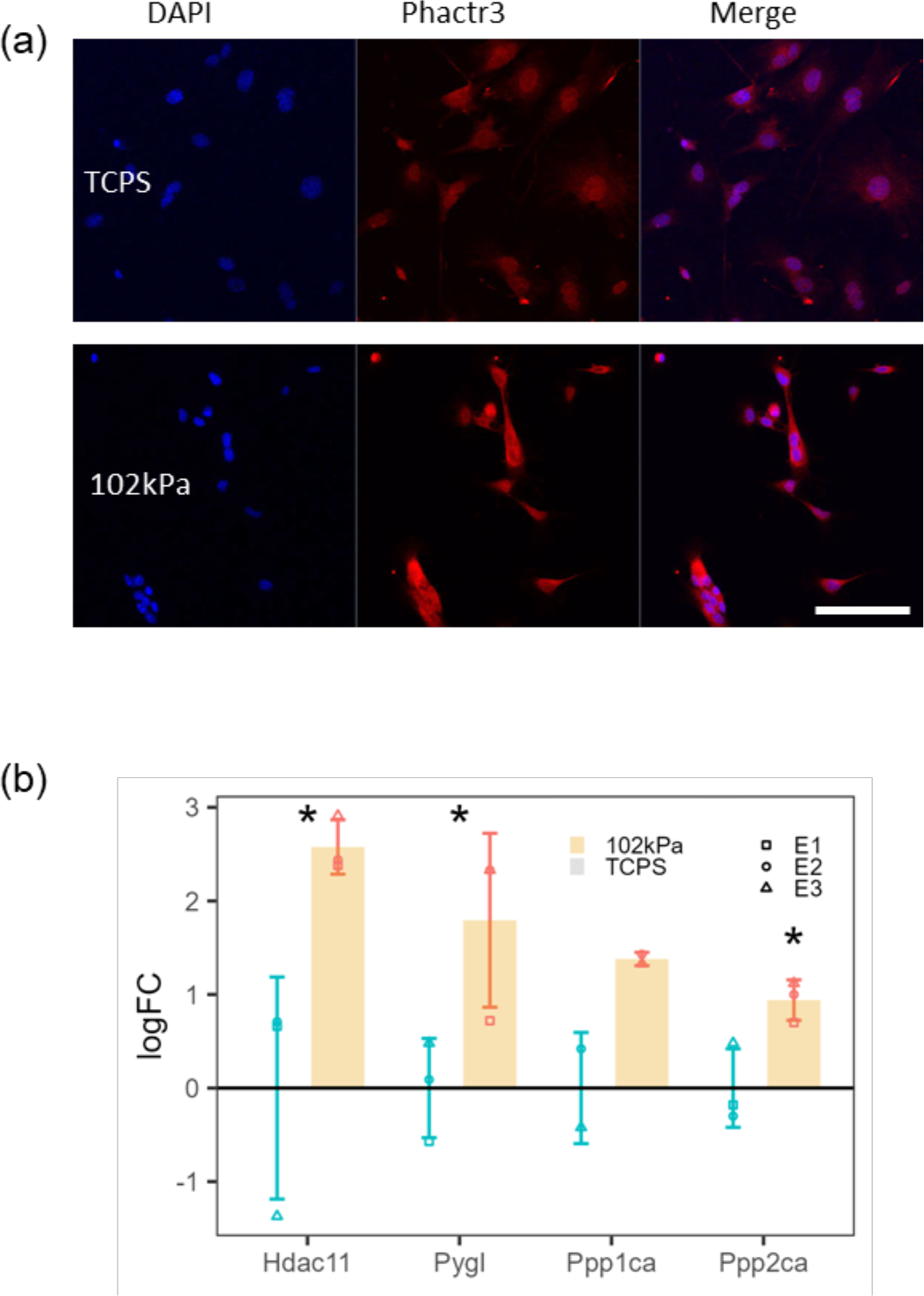
(**a**) Comparison of Phactr3 expression between 102 kPa and TCPS by immunostaining after overnight incubation. (**b**) Metabolism related gene expression (Log fold change (LogFC) relative to TCPS) in MEF inoculated in TCPS and 102 kPa overnight. Line at y =0 represents mean expression in TCPS. Data is represented as mean ± SD (n = 3) and “***”, “**” and “*” represents *p* < 0.001, *p* < 0.01 and *p* < 0.05, respectively. Scale bar represents 50 µm.

